# Age-dependent structural and morphological changes of the stem cell niche disrupt spatiotemporal regulation of stem cells and drive tissue disintegration

**DOI:** 10.1101/2023.01.15.524122

**Authors:** Michelle A. Urman, Nimmy S. John, ChangHwan Lee

## Abstract

Aging induces a progressive decline in tissue function, which has been attributed to a decrease in stem cell function. A major factor driving this decline is the aging of the stem cell niche but elucidating molecular mechanisms of the niche aging and its effects on stem cell regulation remain a challenge. Here, we use the *Caenorhabditis elegans* distal tip cell (DTC), the mesenchymal niche that employs Notch signaling to regulate germline stem cells (GSCs), as an *in vivo* niche aging model and delineate the molecular details of the DTC/niche aging process and its consequences on GSC function and tissue integrity. Using Notch-dependent transcriptional activation as a direct readout of GSC-DTC/niche interaction and its transcriptional activity as a readout for GSC function, we find that an age-dependent reduction in Notch transcription occurs both at the tissue and the cellular levels, but with its activity at the chromosomal loci remains unaffected. This overall reduction is due to an age-dependent progressive shift in the spatial pattern of Notch-dependent transcription in the germline, resulting in a shift of the GSC pool location and disruption of the tissue integrity. We show that the position of the DTC/niche nucleus determines the location of the Notch-responsive GSC pool, with its correlation to the structure and morphology of the DTC/niche, which also changes during aging. Our findings demonstrate that the stem cell niche undergoes structural and morphological changes during aging and reveal a critical link between these changes and the spatiotemporal regulation of stem cell function.

## Introduction

Aging is characterized by degenerative changes and progressive declines in cellular and tissue functions, eventually leading to death (Harman 1981; Kocsisova et al. 2019). Elucidating the aging process has been a challenge due to its complexity, as aging is a result of a multitude of intrinsic and extrinsic factors interplaying, including DNA damage, accumulation or proliferation of toxic metabolites, and environmental stresses (Oh et al. 2014). Aging occurs at several biological levels, molecular, cellular, and organismal levels, each affecting physiology differently. At the molecular level, aging often induces dysregulation of protein synthesis, misfolding, or conformational changes that can damage protein function or alter protein abundance or localization (Johnson et al. 1999; Mc Auley et al. 2017). At the cellular scale, cells enter senescence or replicative aging that coincides with the buildup of misfolded proteins in the cell, DNA or organelle damages, or misregulation of cellular integrity or polarity (DiLoreto & Murphy 2015). Aging often combines with genetic mutations or health risk factors at the organismal level and increases the susceptibility to age-related diseases such as neurodegeneration, cardiovascular disease, and cancer (Booth & Brunet 2016).

Age-dependent declines in tissue function and integrity have been attributed to a decrease in stem cell function (Brunet et al. 2022; Oh et al. 2014). Indeed, proper and timely regulation of stem cell function is key to tissue development, homeostasis, and regeneration (Charville & Rando 2011; Pan et al. 2007; Rossi et al. 2008). Stem cells can remain activated, which are essentially immortal and rejuvenated over the organismal lifecycle, or become quiescent upon the completion of development, exhibiting a progressive decline in their functions and frequency, which has been considered to be aging (Charville & Rando 2011; Devaraj et al. 2021; Pan et al. 2007; Rossi et al. 2008). However, recent studies suggest that stem cells age very little intrinsically, and extrinsic factors such as the microenvironment are largely responsible for the age-dependent stem cell decline (Kalamakis et al. 2019; Morrow & Moore 2019). Unraveling the molecular mechanisms of age-dependent stem cell decline is crucial to advancing therapeutics to delay tissue deterioration, reverse the aging process, and alleviate age-related diseases.

A major extrinsic factor for stem cell function is the stem cell niche, which is located adjacent to stem cells and provides a microenvironment to them. The niches are often specialized, somatic cells and hence are not rejuvenated but experience the aging process, including a structural or functional decline (Jasper & Kennedy 2012; Morrow & Moore 2019; Ryu et al. 2006; Wallenfang 2007). For example, the stem cell niche in the female *Drosophila* gonad progressively loses physical contact with the stem cells as it ages (Pan et al. 2007), and the niche signaling from the neural stem cell niche decreases during aging, inhibiting the stem cells from exiting the quiescent state to repopulate upon injury (Kalamakis et al. 2019; Morrow & Moore 2019). Niche aging has been thought to be a major driver of age-dependent stem cell loss and its functional decline (Brunet et al. 2022; Goodell & Rando 2015; Navarro Negredo et al. 2020). However, molecular mechanisms of the niche aging process and how it affects stem cell physiology and tissue homeostasis remain poorly understood.

The niches regulate stem cells using intercellular niche signaling pathways such as Notch signaling, Wnt signaling, and Hedgehog signaling, which are often highly contextual (Wang et al. 2009). The timing and duration of the signaling transduction and the pattern of its target gene expression vary depending on contexts, such as the tissue type, developmental stage, and environmental conditions (Carlson et al. 2008; Housden & Perrimon 2014). Therefore, analyzing niche signaling in its native contexts is crucial to the precise understanding of *in vivo* niche regulation on stem cells and their aging process. Here, we use the *Caenorhabditis elegans* gonad as an *in vivo* niche aging model, where the mesenchymal distal tip cell (DTC) serves as a niche for a pool of 35-70 germline stem cells (GSCs) (Crittenden et al. 1994; Crittenden et al. 2006; Kimble & Crittenden 2007; Lee et al. 2016) (Figure 1A). The DTC/niche, located at the distal end of the gonad, is essential for germline tissue organization, GSC regulation, and gametogenesis (Austin & Kimble 1987; Byrd et al. 2014; Byrd & Kimble 2009; Cinquin et al. 2010; Crittenden et al. 1994; Kimble & Crittenden 2007; Kimble & White 1981). The DTC/niche is composed of a cap with elaborate cellular processes that enwrap the distal gonad and infiltrate into the GSC pool to ensure physical interactions between the DTC/niche and GSCs for the niche signaling transduction (Linden et al. 2017; Wong et al. 2013). The *C. elegans* germline is a uniquely powerful model to study *in vivo* stem cell-niche interactions and their aging processes. The DTC/niche and GSCs have been well characterized, and their interaction relies on one niche signaling, the Notch signaling pathway, which is continuous throughout the life of the animal (Henderson et al. 1994; Kimble & Seidel 2008; Lee et al. 2016). The simplicity and transparency of *C. elegans* allow for high-resolution whole-tissue or whole-animal imaging (Lee et al. 2016; Lee et al. 2019). In addition, *C. elegans* exhibits an age-dependent decline similar to humans in their physiology and biological functions, including tissue degradation and reproductive decline (Zhang et al. 2020).

**Figure 1:**
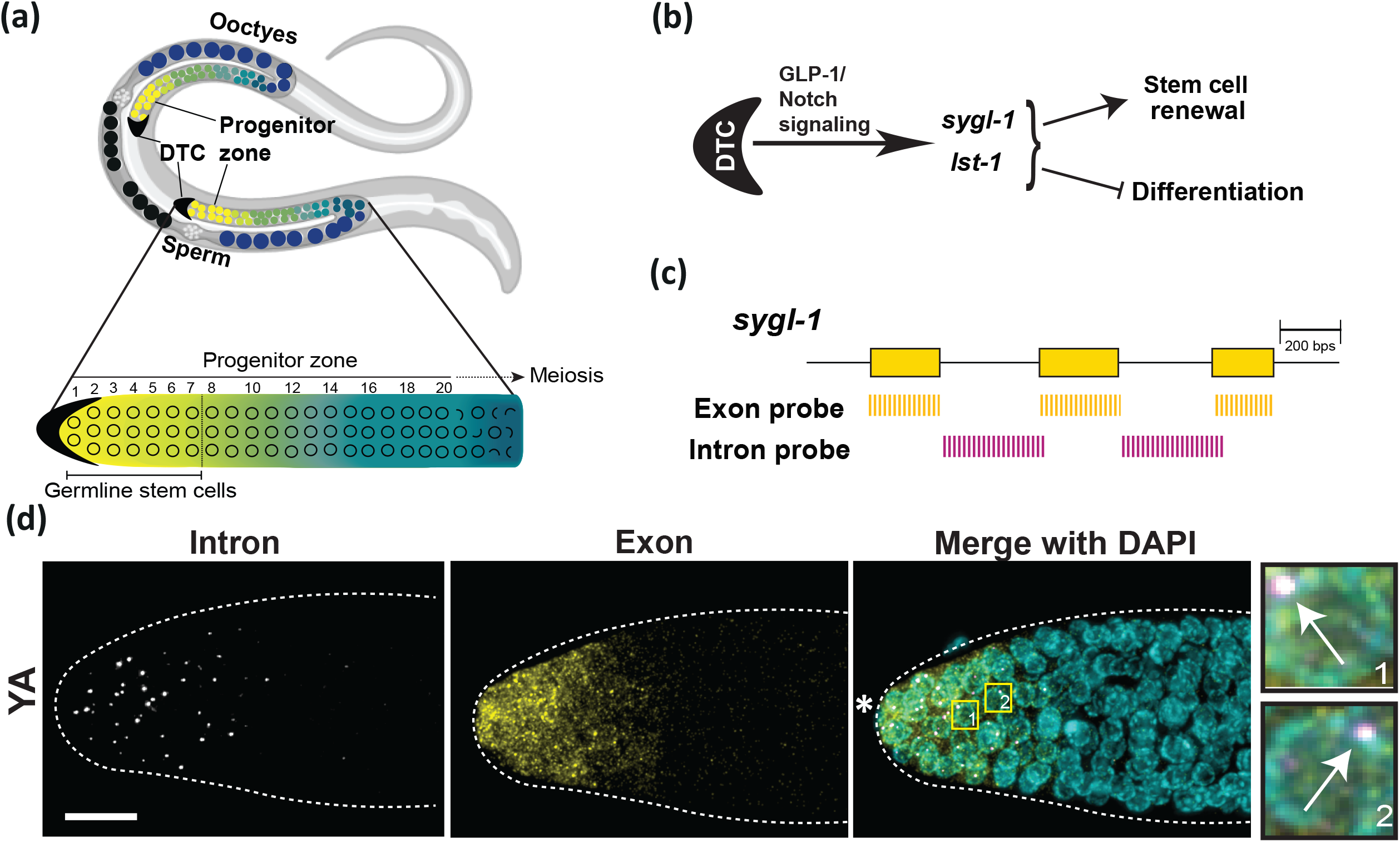
Notch signaling activates transcription of its targets, *sygl-1* and *lst-1* to maintain a germline stem cell (GSC) pool in the *C. elegans* gonad. (a) Schematic of adult *C. elegans* hermaphrodite with U-shaped gonads with a distal tip cell (DTC) at each distal end (black crescents). Along the proximal end, oocytes (dark blue) reside that meet with sperm to form eggs (black). Germline stem cells (GSCs; yellow) occupy the first 6-8 germ cell rows of the germline. The cells progress through the progenitor zone, and cells enter differentiation (green) and enter meiosis (blue) before becoming oocytes. (b) The distal tip cell (DTC/niche) employs GLP-1/Notch signaling to activate transcription from the direct downstream target genes, *sygl-1*, and *lst-1*, to maintain the germline stem pool by promoting self-renewal and preventing differentiation. (c) A schematic of the smFISH probes used in this study. The design of two smFISH probe sets used to visualize *sygl-1* transcripts. D. Z-projection of *sygl-1* smFISH signal in the distal gonad in the wild-type young adult (YA, 24 h post mid-L4). The intron probes (Left) indicate only nascent transcripts whereas the exon probes (middle) show both the nascent transcripts (bright dots) and cytoplasmic mRNAs (small, dim dots). DAPI marks nuclei (right). The small white boxes (right) are 10X zoomed in on the right. The bright white dots overlapping with DAPI (arrows) confirm *sygl-1* active transcription sites (ATS). White asterisks indicate the position of the DTC nuclei. Scale bar: 10 μm.

The DTC/niche uses Notch signaling to regulate GSCs in the *C. elegans* gonad (Byrd et al. 2014; Lee et al. 2016; Byrd & Kimble 2009). This broadly conserved cell-cell signaling pathway is activated throughout metazoan for important biological functions such as cell fate decision, cell proliferation, apoptosis, and tissue patterning, with its dysfunction leading to various diseases including cardiovascular disease and cancer (Bocci et al. 2020; Kopan 2012; Siebel & Lendahl 2017). Notch is activated upon ligand-receptor interaction, which triggers a series of proteolytic cleavages on the Notch receptor (GLP-1 or LIN-12 in *C. elegans*) and releases the Notch intracellular domain (NICD) that translocates into the nucleus and activates transcription from targets with a DNA-binding protein, CSL/RBPJ-κ, and a co-activator, MAML (LAG-1 and LAG-3/SEL-8 in *C. elegans*, respectively) (Bray 2006; Chen et al. 2020; Kershner et al. 2014). In the *C. elegans* gonad, GLP-1/Notch activates at least two well-studied target genes, *sygl-1*, and *lst-1*, both of which encode stem cell effectors (Figure 1B) (Brenner & Schedl 2016; Chen et al. 2020; Kershner et al. 2014; Lee et al. 2016; Singh et al. 2011). Nascent transcripts from the active transcription sites (ATS) of the two targets have been used as direct readouts of Notch activation, which revealed a stochastic but probabilistic nature of Notch signaling (Crittenden et al. 2019; Lee et al. 2016; Lee et al. 2019; Lynch et al. 2022). Importantly, the distribution of *sygl-1* ATS coincides with the GSC pool, thus has been established as a potent marker for the GSC pool (Cinquin et al. 2010; Crittenden et al. 2019; Kershner et al. 2014; Lee et al. 2016; Lee et al. 2019; Shin et al. 2017). Despite extensive studies delineating Notch functions for the niche-stem cell interactions, little is known about the link between niche aging and Notch activation.

Here, we analyze the DTC/niche aging process and its effects on Notch activation and GSC function in aging *C. elegans*, from its young adult stage (YA or 24h post the mid-L4 larval stage) to Day 3 since YA (A72 or 72h post-YA), where the worm exhausts its premade sperm supply and concludes self-fertility (Andux & Ellis 2008; Byerly et al. 1976; Kocsisova et al. 2019; Scharf et al. 2021). We observe an age-dependent decline in Notch-dependent transcription both at the tissue level and the cellular level, even on Day 1 of aging. However, the Notch targets sustain the same transcriptional activity at their individual chromosomal loci through aging. We find that the overall reduction in Notch activation stems from an age-dependent progressive shift in the spatial pattern of Notch-dependent transcription, which in turn dislocates the GSC pool in the distal gonad and disrupts the tissue polarity. Finally, we reveal a strong correlation between Notch activation and the position of the DTC/niche nucleus, which also gradually shifts with aging, and show their link to age-dependent changes in the structure and morphology of the DTC/niche.

## Results

### Aging process begins early in adulthood and induces a decline in Notch-dependent transcriptional activation, leading to a reduction of the GSC pool

To elucidate mechanisms of niche aging and its effects on stem cells *in vivo*, this study focuses on the *C. elegans* gonad, where a single mesenchymal cell, the distal tip cell (DTC) uses GLP-1/Notch signaling to maintain a pool of 35-70 germline stem cells (GSCs) at the distal gonad under normal conditions (Cinquin et al. 2010; Crittenden et al. 2017) (Figure 1A-B). To precisely localize the GSCs and examine their function, we used well-established Notch readouts, the active transcription sites (ATS) and mRNAs from a Notch target, *sygl-1*, and directly measured Notch-dependent transcriptional activation, which has been used as a barometer for GSC function and stemness (Austin & Kimble 1987; Brenner & Schedl 2016; Kimble & White 1981; Lee et al. 2016; Sorensen et al. 2020) (Figure 1B-D). Briefly, we used single-molecule fluorescence *in situ* hybridization (smFISH) with two probe sets, one targeting *sygl-1* introns and the other targeting exons (Figure 1C), which revealed both the nuclear nascent transcripts at *sygl-1* ATS and cytoplasmic *sygl-1* mRNAs at the single-molecule level (Figure 1D) (Lee et al. 2016; Raj & van Oudenaarden 2009). The smFISH results allow for precise quantification of transcription, such as its activation rate and the level of its activity in the whole germline tissue, in each GSC, and at each *sygl-1* chromosomal locus, through aging (Lee et al. 2016; Lee et al. 2017; Raj & van Oudenaarden 2009).

Aged *C. elegans* exhibits organismal aging phenotypes, including declines in locomotion, egg-laying rate, and germ cell division, along with GSC quiescence (Byerly et al. 1976; Collins et al. 2008; Scharf et al. 2021; Seidel & Kimble 2015; Zhang et al. 2020). Consistently, the size of the progenitor zone (PZ) in the germline, which harbors the mitotic germ cells including a GSC pool, also decreased progressively during aging (Kocsisova et al. 2019) (Figure S1A). To analyze the effects of aging on the GSC pool, we performed *sygl-1* smFISH through aging, from the young adult stage (YA or 24h post the mid-L4 stage) to Day 3 since YA (A72 or 72 h post-YA), when there is little to no gametogenesis from self-fertilization (Figure 1D and 2A) and the germ cell division slows down dramatically (Andux & Ellis 2008; Kocsisova et al. 2019; Seidel & Kimble 2015). The results reveal that Notch-dependent transcription remains active in the distal gonad throughout aging, with many *sygl-1* ATS and mRNAs present even at A72 (Figure 2A), which indicates that the DTC/niche-GSC interaction sustains, and Notch is still activated in the aged gonads (Figure 2A). To assess how the GSC pool size changes during aging, we compared the total number of germ cells containing at least one *sygl-1* ATS between young and aged worms (Figure 2B). The number of *sygl-1* ATS-containing cells generally declined, with the most aged group (A72) exhibiting about a 30% decrease compared to YA (Figure 2B). Consistently, the total number of *sygl-1* ATS in the gonad, which estimates the overall Notch-dependent transcriptional activation at the tissue level, also gradually decreased with aging (Figure S1B). However, transcriptional activation of a Notch-independent gene, *let-858* (Lee et al. 2016), remained unchanged during aging, rejecting that aging affected the overall transcription level, at least from YA to A72 (Figure S2). The older worms showed greater variability both in the number of *sygl-1* ATS-containing cells and the total number of *sygl-1* ATS in the distal gonad (Figure 2B and Figure S1B), supporting that Notch regulation is disrupted with aging. Altogether, the results show that Notch-dependent transcriptional activation declines at the tissue level even from the very early stage of aging (A24), accompanied by a reduction of the Notch-responsive GSC pool.

**Figure 2:**
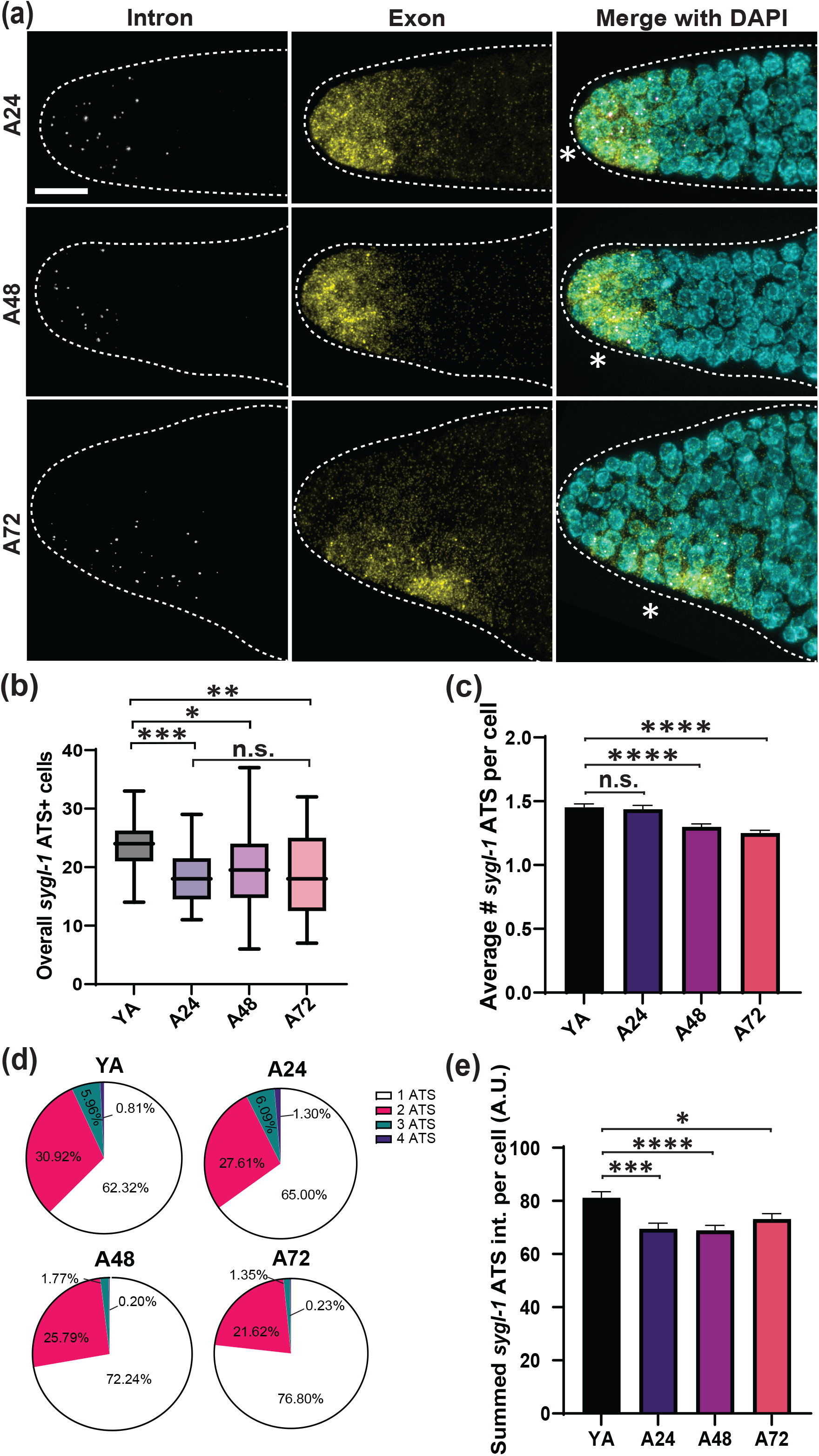
The overall *sygl-1* transcriptional activation decreases with aging. (a) *sygl-1* smFISH of the distal gonad through aging, at 24 h post YA (A24), 48 h post YA (A48), and 72 h post YA (A72). White asterisk: the position of the DTC/niche nucleus. (b) The overall number of *sygl-1* ATS-containing cells per gonad are plotted for each aging stage. For all box-and-whiskers plot in this study, the middle line shows the median; top and bottom of box are the third and first quartiles, respectively; whiskers, maximum and minimum of data points; circles, outliers (value greater than 1.5X first or third quartile from the median). (c) The average number of *sygl-1* ATS per cell is shown for all ages. All error bars in this study are the standard error of the mean (SEM) unless stated otherwise. (d) The GSCs in each gonad are grouped by the number of *sygl-1* ATS that they contain and plotted as fractions for each age. (e) The *sygl-1* ATS intensities are pooled in each GSC and plotted for ages. (b-e) n=27 gonads for YA, n=25 for A24, n=26 for A48, and n=24 for A72. For all t-tests (one sample or two sample) in this study, *p<0.05, **p<0.01, **p<0.001, ***p<0.0001, ****p<0.00001, and *****p<0.000001. ‘n.s.’: not significant by t-test.

### Aging reduces Notch-dependent transcriptional activation in each GSC

Despite a decline in the overall Notch-dependent transcription at the tissue level (Figure 2A-B and Figure S1B), the impact of aging may be different at the cellular level. To test this idea, *sygl-1* transcription was further assessed in the individual GSCs. Each GSC can have up to four *sygl-1* ATS, and a higher ATS number generally indicates a higher Notch activation in that GSC, which tends to occur near the DTC/niche (Austin & Kimble 1987; Kimble & White 1981; Lander et al. 2012). We recorded the number of *sygl-1* ATS in each GSC from YA to A72 and found that the number gradually decreased during aging (Figure 2C), similar to the total number of *sygl-1* ATS-containing cells (Figure 2B). Consistently, the fraction of GSCs containing >1 *sygl-1* ATS progressively declined during aging, with the 4 *sygl-1* ATS-containing GSCs almost completely depleted in A72 (Figure 2D, Figure S3A). The summed *sygl-1* ATS intensity in each GSC, which estimates how many *sygl-1* transcripts are produced and therefore indicates Notch-dependent transcriptional activity of the GSC, also declined with aging (Figure 2E and Figure S1C). We conclude that aging occurs both at the tissue (Figure 2B) and the cellular levels (Figure 2C-E), showing reduction both in Notch-dependent transcriptional activation and its activity in the GSC.

### Aging drives progressive changes in the spatial pattern of Notch-dependent transcription in the gonad

Notch-dependent transcription is steeply graded within the GSC pool, with its probability and activity highest next to the DTC/niche under normal conditions (Crittenden et al. 2019; Lee et al. 2016; Lynch et al. 2022). This graded spatial pattern of Notch activation is crucial for regulating the size and function of the GSC pool (Crittenden et al. 2019; Lee et al. 2016; Lynch et al. 2022). Mutations in the Notch pathway often modulate this spatial pattern, leading to pathologies such as tumorigenesis and stem cell loss (Izumchenko et al. 2015; Kaylan et al. 2018; Lee et al. 2016; Mishra et al. 2001). Therefore, maintaining the ‘normal’ gradient of Notch activation is key to tissue homeostasis and functionality. We hypothesized that aging might induce changes in the spatial pattern of Notch-dependent transcriptional activation, leading to disruption of the GSC pool. To analyze the spatial pattern of Notch transcription, we recorded the percentage of germ cells containing at least one *sygl-1* ATS as a function of distance from the distal end through aging, which estimates the probability of Notch transcriptional activation at various positions relative to the DTC/niche (Figure 3A-B). The YA worms showed the ‘normal’ spatial pattern for *sygl-1* transcription as expected, exhibiting a steep gradient within the GSC pool (0-25 μm from the distal end of the gonad), from about 70% at the distal-most region to less than 2.5% at the border of the GSC pool (Figure 3A, YA) (Lee et al. 2016). The graded pattern was maintained through aging (Figure 3A). However, the percentage of *sygl-1* ATS-containing cells was steadily reduced at the distal gonad as the worm aged, flattening the gradation (Figure 3A and Figure S3A, 0-15 μm from the distal end). Notably, the proximal pool of GSCs gradually increased during aging (Figure 3A, 15-35 μm from the distal end), extending the GSC boundary more proximally (Figure 3A, red dashed lines).

**Figure 3:**
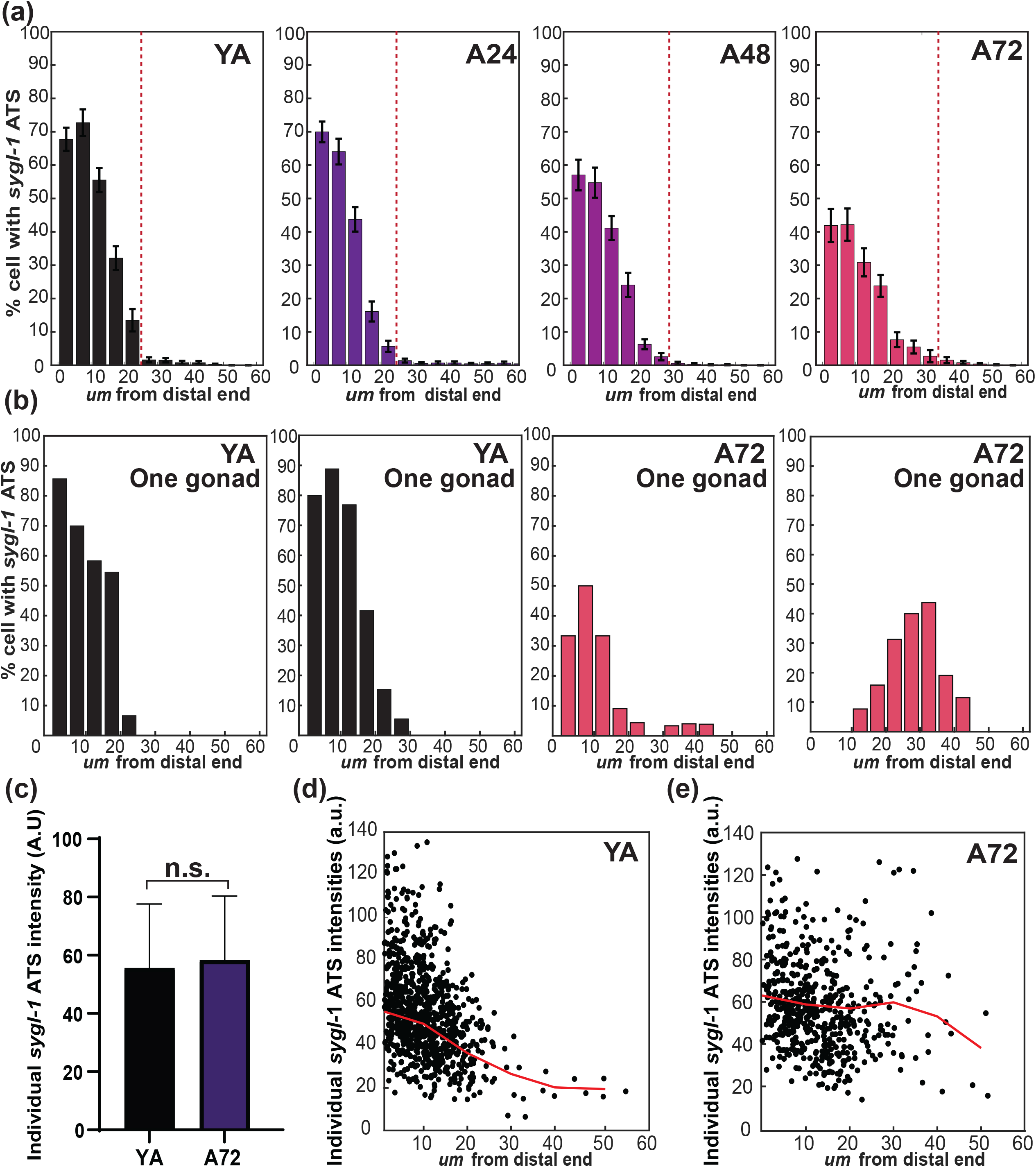
Aging induces a progressive shift in the spatial pattern of Notch activation. (a) The percentage of cells containing at least 1 *sygl-1* ATS was plotted as a function of the distance from the distal end of the gonad for each age. Red dashed line indicates where the percentage is lower than 2.5%, which marks the boundary of the GSC pool. n=27 gonads for YA, n=25 for A24, n=26 for A48, and n=24 for A72. (b) Examples of the individual spatial pattern of *sygl-1* transcriptional activation in one gonad at YA (black) or A72 (magenta), separated from the average spatial pattern in (a). (c) The individual *sygl-1* ATS intensities are compared between YA and A72. (d-e) The Individual *sygl-1* ATS intensities are plotted against the distance from the distal end in YA (d) or A72 (E). Each dot represents one *sygl-1* ATS. (c-e) n= 931 ATS for YA and 550 for A72.

To understand how aging increased Notch activation in the proximal pool of GSCs, we examined *sygl-1* transcriptional patterns in the individual gonads (Figure 3B), which could reveal details masked in the average spatial pattern (Figure 3A). The YA worms showed individual spatial patterns similar to the average pattern with small variability (compare Figure 3B to Figure 3A; black plots). In most YA gonads, the percentage of *sygl-1* ATS-containing cells peaked generally at the distal-most region of the gonad with a steep gradient that ends at around 25 μm (Figure 3B, YA). In A72 gonads, however, the graded pattern became less obvious, and its peak was shifted farther from the distal end with relatively higher variability in its position (Figure 3B, A72). Many A72 gonads completely lost *sygl-1* ATS-containing cells at the distal gonad (Figure 3B, A72). Also, several aged gonads exhibited premature GSC differentiation at the distal gonad, containing cells in meiotic prophase (Figure S3B). We conclude that aging disrupts the ‘normal’ spatial pattern of Notch-dependent transcription, shifting its peak to a proximal gonad and disrupting germline tissue polarity.

### Transcriptional activity on each *sygl-1* chromosomal locus is unaffected by aging

Despite the overall decrease and locational shift of Notch-dependent transcriptional activation in the gonad during aging, the individual *sygl-1* ATS intensities, which estimate transcriptional activity on each *sygl-1* chromosomal locus, remained unchanged in the aged worms (Figure 3C and Figure S3C). The individual *sygl-1* ATS intensities also showed a graded pattern in YA (Figure 3D), but the pattern lost the gradient in A72 (Figure 3D-E and Figure S3C), reflecting the age-dependent changes in *sygl-1* transcriptional activation (Figure 3A-B). Altogether, these results show that aging leads to declines in the overall Notch activation at the tissue and cellular levels but not at the chromosomal level.

### The abundance and spatial pattern of *sygl-1* mRNAs are minimally impacted by aging

Another well-established readout for Notch activation is *sygl-1* mRNA, which has been used complementarily to assess the GSC function (Crittenden et al. 2017; Lee et al. 2016; Lynch et al. 2022; Shin et al. 2017). To analyze the spatial pattern of *sygl-1* mRNAs, we recorded the number of *sygl-1* mRNAs per cell (Figure S4A) or the percentage of cells containing *sygl-1* mRNAs (Figure S4B) as a function of distance from the distal end. The spatial patterns of *sygl-1* mRNAs from both assays exhibited a gradient with the mRNA counts highest at the distal-most gonad (Figure S4A), resembling the pattern of *sygl-1* ATS-containing cells in YA (Figure 3A). However, the mRNA abundance was generally comparable in all ages at the corresponding positions within the GSC pool (Figure S4A). Notably, there was a slight increase in the proximal pool of GSCs in A72 (Figure S4A, A72, >30μm), consistent with an increase in *sygl-1* transcriptional activation in that region at A72 (Figure 3A, A72). The percentage of cells containing *sygl-1* mRNAs showed essentially the same trend as the mRNA number per cell, with almost all germ cells located at the distal region containing the mRNAs (Figure S4B). There was an increase in the percentage of cells containing *sygl-1* mRNAs at the proximal region in A72 (Figure S4B, A72, >30 μm), which is likely due to the Notch-independent *sygl-1* expression during meiotic prophase, another indication of a shorter PZ in the aged gonads (Figure S4B) (Kershner et al. 2014; Lee et al. 2016).

### The DTC/niche nucleus gradually shifts to the proximal gonad along with the Notch-responsive GSC pool during aging

What causes the shift of Notch activation to a proximal region of the gonad during aging? Is the shift related to the location, function, or morphology of the DTC/niche, or is it merely from stochasticity? To answer these questions, we examined the cellular features of the aging DTC/niche (Figure 4). The DTC/niche is a mononuclear mesenchymal cell with its nucleus typically located at the distal tip of the gonad through development and at YA (Kimble & White, 1981) (Figure 4A and Figure S5A, YA). However, the DAPI staining showed that the DTC/niche nucleus no longer remained at the distal tip but was located at a more proximal region in the majority of A72 gonads (Figure 4A and Figure S5A). Indeed, the DTC/niche nucleus progressively shifted from the distal-most end of the gonad (∼2 μm from the distal tip) at YA to a proximal region (∼12 μm from the distal tip) at A72, at least 5 times farther from the distal tip of the gonad compared to YA, with a maximum shift over 22 μm (Figure 4B-C). The variability of the DTC/niche nucleus shift also increased with aging, suggesting that regulation holding the DTC/niche nucleus in the original place might be weakened with aging (Figure 4C).

**Figure 4:**
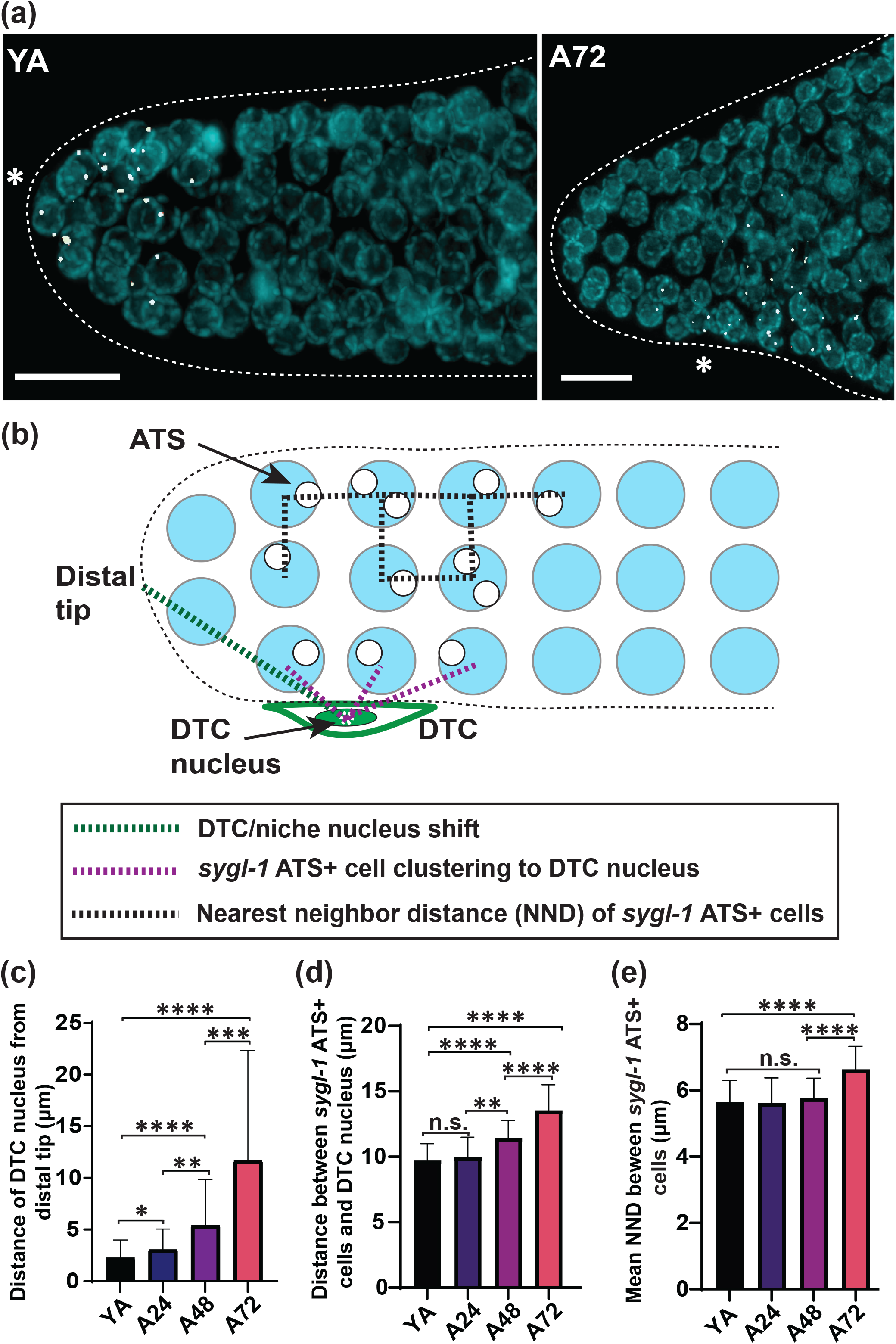
*sygl-1* ATS-containing cells are clustered to each other and to the DTC/niche nucleus. (a) Representative *sygl-1* smFISH with DAPI staining in YA (left) and A72 (right). The asterisk marks the location of the DTC/niche nucleus. Scale bar: 10 μm. (b) A schematic of the spatial analyses of Notch activation used in (c-e). (c) The Euclidean distances between the DTC/niche nucleus and the distal tip of the gonad, indicating the amount of nuclear shift, are measured through aging. n=45 gonads for all ages. Error bar: the standard deviation. (d) The distance between the sygl-1 ATS-containing cells and the DTC/niche nucleus to estimate the clustering of Notch activation to the nucleus through aging. (e) The shortest distance between two neighboring cells that contain at least 1 *sygl-1* ATS (the nearest neighbor distance, NND) are shown for all ages. (d-e) n=27, 25, 26, and 24 gonads for YA, A24, A48, and A72, respectively.

Given the similar proximal shift both in the *sygl-1* ATS-containing cells (Figure 3A-B) and the DTC/niche nucleus (Figure 4C), we wondered if there is a link between them. For example, *sygl-1* ATS may appear only near the DTC/niche nucleus, and an age-dependent shift of the DTC/niche nucleus may move *sygl-1* ATS as well, keeping them nearby. To test this idea, we measured how clustered the *sygl-1* ATS-containing cells were to the DTC/niche nucleus, by recording the distances between the DTC/niche and all *sygl-1* ATS-containing cells in each gonad from YA to A72 (Figure 4D). The average distance was about 10 μm at YA, about half the length of the GSC pool, and only slightly increased to 13 μm in A72, less than one germ cell diameter (Lee et al. 2016) (Figure 4D). These results indicate that Notch activation occurs mostly near the DTC/niche nucleus and this clustering remains in the aged gonads. Furthermore, the nearest neighbor distances (NNDs) of *sygl-1* ATS-containing cells in each gonad, which estimate the degree of clustering within the Notch-responsive GSCs, showed little to no changes during aging (Figure 4E). We also used *sygl-1* mRNAs instead of ATS for similar assays, which showed essentially the same results, further supporting the idea of sustained Notch activation near the DTC/niche nucleus regardless of age (Figure S5D-E). Consistently, the size of the region where the *sygl-1* ATS-containing cells are clustered remains unchanged during aging as well as the region where *sygl-1* mRNAs are rich (Figure S5B-C), despite their locational changes due to aging (Figure 3A-B). A72 exhibited greater variability overall, reflecting the high variability in the DTC/niche nucleus shift at the age (Figure 4C). We conclude that Notch activation occurs near the DTC/niche nucleus, keeping GSCs clustered even when they shift proximally in the aged gonads.

### The distance to the DTC/niche nucleus determines Notch-dependent transcriptional activity in the GSC

To further investigate the link between the DTC/niche nucleus and Notch activation, we examined the summed *sygl-1* ATS intensities in the individual GSCs, estimating Notch-dependent transcriptional activity per cell, with their positions relative to the DTC/niche nucleus (Figure 5A and Figure S6A). The analysis revealed that there is a modest correlation between the *sygl-1* transcriptional activity in a GSC and the proximity of the GSC to the DTC/niche nucleus in YA. This correlation remained through aging, although the correlation became slightly weaker at the older stages (Figure 5A, A48 and A72). We also used a second Notch readout, *sygl-1* mRNAs for a similar analysis, which showed much stronger correlations between the number of *sygl-1* mRNAs in a cell and the proximity to the DTC/niche nucleus, throughout aging (Figure 5B), albeit a slight decrease of the correlation in more aged gonads, similar to the summed *sygl-1* ATS intensities (Figure 5A). The same analysis using the individual gonads also showed a very similar correlation and the same age-dependent effects (Figure S6). Altogether, we find a strong correlation between Notch transcriptional activity in the GSC and its distance to the DTC/niche nucleus, and this correlation remains during aging albeit with a slight decrease. The results suggest that the proximity to the DTC/niche nucleus determines how active Notch-dependent transcription will be.

**Figure 5:**
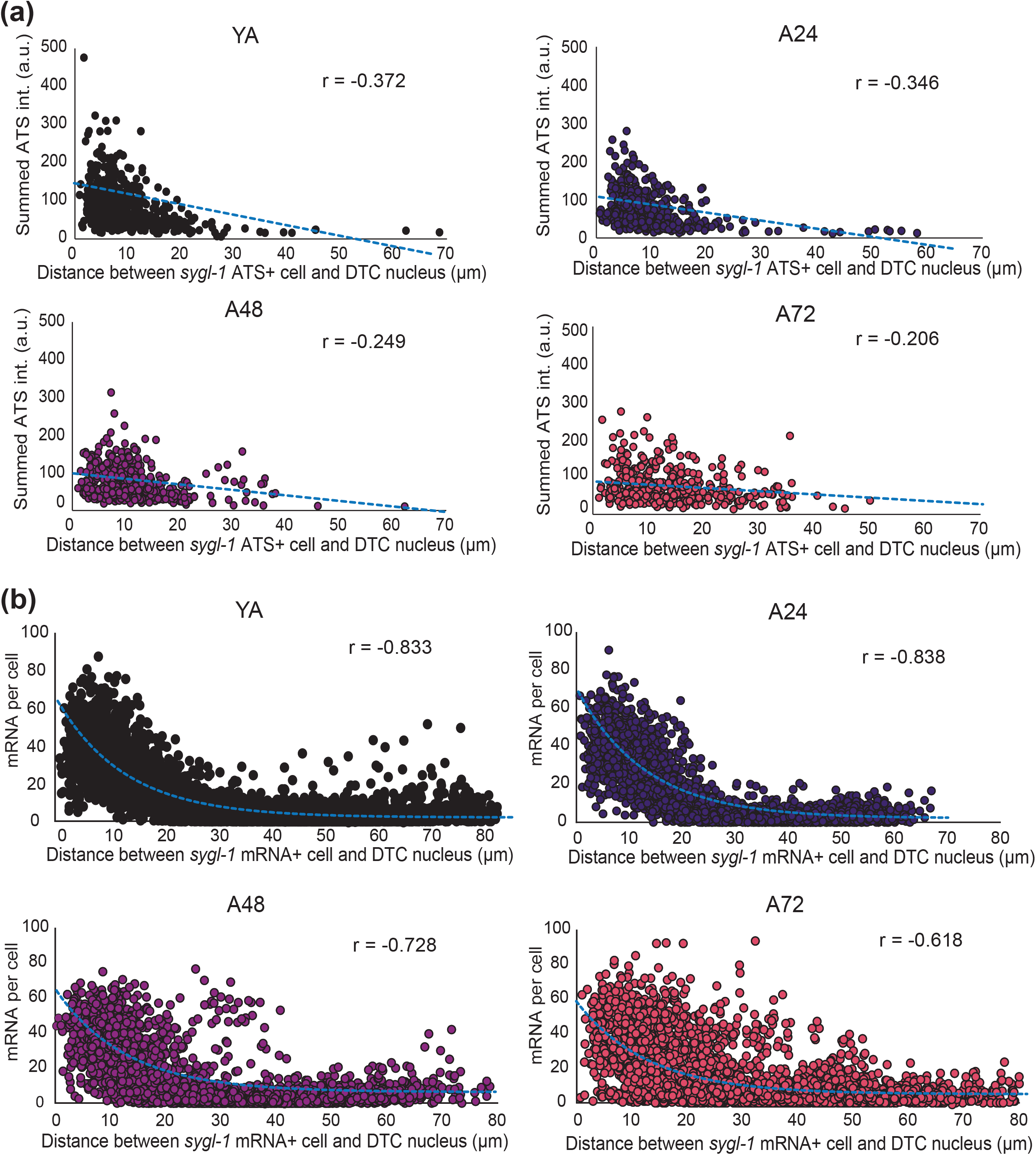
*sygl-1* transcription is activated near the DTC/niche nucleus, with a correlation between Notch transcriptional activity and the proximity to the nucleus. (a) The summed *sygl-1* ATS intensities in each cell are plotted against the distance between the corresponding cell and the DTC/niche nucleus for all ages. ‘r’ indicates the Pearson’s correlation coefficient. n=621, 620, 508, and 444 nuclei for YA, A24, A48, and A72, respectively. (b) Total number of mRNAs per cell as a function of the distance between the corresponding cell and the DTC/niche nucleus. n=5,462, 3,913, 3,914, and 3,914 cells for YA, A24, A48, and A72, respectively. (a-b) The blue dashed line indicates the line fitting model to calculate Pearson’s r value.

### Aging affects the structure and integrity of the DTC/niche

The *C. elegans* DTC/niche has an extensive structure, consisting of a cap that encapsulates a distal half of the GSC pool and the cellular processes, the tubular structures, that extend from the cap and reach the germ cells even beyond the GSC pool (Byrd et al. 2014; Crittenden et al. 2006; Wong et al. 2013). The processes fall into two groups, the long-extended processes (LEPs) that are typically thick and long to reach the end of the progenitor zone (PZ), and the short intercalating processes (SIPs) that infiltrate into the gonad to physically interact with the germ cells mostly at the distal gonad (Byrd & Kimble 2009; Byrd et al. 2014). Visualization of the DTC/niche plasma membrane using the myristoylated GFP (myrGFP) expressed under a DTC/niche specific promoter, *lag-2*, revealed the cellular structure of the DTC/niche during aging, including the cap, LEPs, and SIPs (Figure 6A). The length of LEPs increased during aging, with LEPs in A72 about 30% longer than YA, similar to the previous reports (Crittenden et al. 2006) (Figure S7A). However, there was little increase in the SIP extent from YA to A72 (Figure 6B), suggesting little to no change in DTC/niche-GSC connectivity during aging. The cap of the DTC/niche has a few gaps, where the myrGFP signal is missing and therefore the DTC/niche cannot make physical contact with the germ cells for Notch activation (Byrd et al. 2014; Crittenden et al. 2006; Lee et al. 2016; Byrd & Kimble 2009) (Figure S7B). The DTC/niche gaps grew larger during aging by over two folds from YA to A72, showing that aging disrupts the cellular integrity of the DTC/niche (Figure 6C and Figure S7B). However, the total number of DTC/niche gaps did not exhibit any age-dependent change (Figure 6D). In addition, the total number of cellular fragments detached from the DTC/niche decreased significantly from YA to A72 (Figure 6E), suggesting that aging affects the DTC/niche dynamics in forming new processes or severing newly formed processes, which indeed occur frequently under normal conditions (Wong et al. 2013). The size of the detached DTC fragments decreased in size as well at the older stages (Figure S7C, A48 and A72). These results show that the DTC/niche undergoes structural and morphological changes during aging, which coincide with the shift of the DTC/niche nucleus and age-dependent changes in the spatiotemporal pattern of Notch activation, which in turn lead to a decline in the GSC function and disruption in the germline tissue integrity.

**Figure 6:**
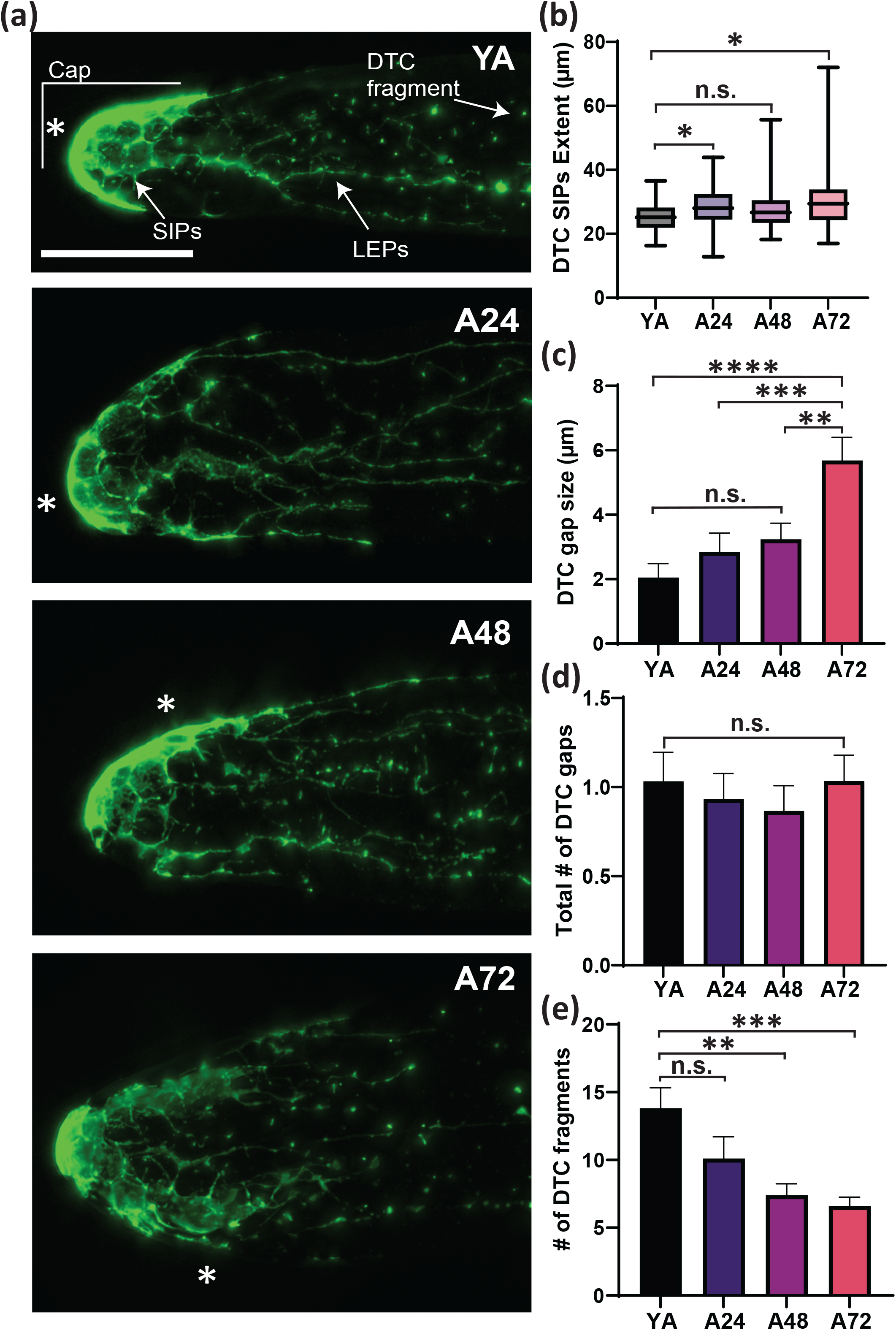
Aging induces changes in the structure and morphology of the DTC/niche. (a) Myristoylated GFP expressed under the DTC-specific promoter, *lag-2*, visualizes the plasma membrane of the DTC/niche. The maximum Z projection is shown. Asterisk marks the location of the DTC/niche nucleus. LEPs: long external processes. SIPs: short intercalating processes. Scale bar: 20 μm. (b) The distance between the distal tip and the most proximally located SIP is plotted, estimating the SIP extent in the gonad in all ages. n=45 for all ages. (c) The size of the DTC gap, a region missing its plasma membrane for all ages. (d) The total number of DTC gaps in a gonad for all ages. (c-d) n=30, 30, 30, and 29 gonads for YA, A24, A48, and A72, respectively. (e) The number of detached DTC fragments, isolated small DTC/niche membranes. n=10 gonads for all ages.

## Discussion

This work analyzes the *in vivo* stem cell niche aging process and its consequences at various biological levels, the organismal, tissue, cellular, and chromosomal levels, using the aging *C. elegans* gonad with a focus on the DTC/niche-GSC interactions that are mediated by Notch signaling. This study examines Notch-regulated transcriptional activation and its activity, which are well-established sensitive and accurate metrics for Notch activation (Lee et al. 2016; Lee et al. 2017; Lynch et al. 2022), from the young adult stage (YA), where the hermaphrodite worm becomes gravid, to Day 3 since YA (A72), when the worm concludes self-fertilization and ceases embryogenesis without the external sperm from mating (Scharf et al. 2021; Van Bael et al. 2018). We observe dramatic changes in Notch-dependent transcription in the GSCs even at the early stage of aging (A24), including an overall decline in Notch activation and a progressive shift in the spatial pattern and location of Notch-dependent transcription, which leads to a shrinkage and locational shift of the GSC pool within the germline. We show that these changes are due to an age-dependent shift of the DTC/niche nucleus and reveal a strong correlation between Notch-dependent transcription and its proximity to the DTC/niche nucleus. We also find that this relationship is linked to age-dependent structural and morphological changes in the DTC/niche. Below we place the gained insights in context and discuss their broader implications.

### The DTC/niche aging effects are specific to Notch-dependent transcriptional activation

One of the major findings in this study is that aging reduces overall Notch-dependent transcriptional activation in the whole germline and within each GSC, whereas there is no change in transcriptional activity, the number of nascent transcripts synthesized at each *sygl-1* locus during aging (Figure 2 and Figure 3C-E). This means that age-dependent declines occur both at the tissue and the cellular levels, but not at the chromosomal level. Our findings are consistent with the previous studies showing age-dependent tissue-level declines (Crittenden et al. 2006; Kocsisova et al. 2019; Scharf et al. 2021), but using our direct Notch readouts allowed us to further analyze stem cell-niche interactions and Notch activation at various biological levels with single-molecule precision. At the tissue level, aging decreases the number of germ cells responding to Notch signaling, causing a reduction in the Notch-responsive GSC pool (Figure 2A-C and Figure S1). At the cellular level, aging reduces the number of *sygl-1* ATS per cell, meaning that the probability of Notch-induced transcriptional firing in the GSC decreases by aging, which indicates a decline in GSC function (Figure 2D-E). At the chromosomal level, however, the number of transcripts synthesized at each *sygl-1* locus, estimated by the *sygl-1* ATS intensity, remains unaffected by aging (Figure 3C-E). Transcription of a Notch-independent gene, *let-858*, also remains unchanged (Figure S2), suggesting that the aging effects are specific to Notch activation from the DTC/niche and its responses in the GSCs. The capacity or function of transcriptional machinery in the germ cells including GSCs is not altered during aging, supporting the idea that the age-dependent stem cell decline is not due to the aging of the stem cells *per se* but rather a consequence of the niche aging and its functional decline (Kalamakis et al. 2019; Morrow & Moore 2019; Navarro Negredo et al. 2020; Oh et al. 2014). Examining a second Notch target, *lst-1* will further confirm the conclusion. Notably, the DTC/niche aging occurs early in adulthood, as early as Day 1 since YA, similar to the aging of some human tissues that begins soon after puberty, such as connective tissue of the eye and skin (Freemont & Hoyland 2007; Pardue & Sivak 2000). In general, older gonads exhibit higher variability in Notch-dependent transcription, suggesting that the age-dependent declines may be in part due to the regulation of DTC/niche becoming weak and inconsistent or GSCs becoming more susceptible to gene expression noise of Notch targets, possibly leading to inefficient self-renewal and early differentiation. Analyzing the aging process and its effects on other tissues or organisms and expanding the assays to even older stages will broaden our current understanding.

### The DTC/niche nucleus defines the Notch-responsive GSC pool

We show that there is a progressive change in the spatial pattern of Notch-activated transcription, from a steep probability gradient at YA to a more flattened gradient both in *sygl-1* ATS occurrence and mRNA abundance. During aging the probability of Notch activation decreases at the distal-most gonad, where it is normally highest at YA, whereas the probability at a more proximal region increases over time (Figure 3 and S4). These results contradict the overall reduction of the Notch-responsive GSC pool (Figure 2) as these changes seemingly extend the GSC pool at the proximal region (Figure 3A). We find that the change in the spatial pattern is due to the shift of the location of Notch activation to a proximal gonad, leaving little to no Notch-responsive GSCs at the distal gonad, rather than an expansion of the GSC pool (Figure 3B). As a result, the crescent cells transitioning to gametes, which are germ cells in the meiotic prophase, emerge at the distal region of the aged gonads, disrupting the germline tissue integrity and polarity (Figure S3B).

Then, what drives this locational shift of Notch activation? Strikingly, we find a correlation between Notch-dependent transcriptional activity in a cell and its proximity to the DTC/niche nucleus, and this correlation remains through aging albeit a slight decrease in the older gonads (Figure 5A), consistent with a slight decrease in clustering of *sygl-1* ATS-containing cells to the DTC/niche nucleus (Figure 4D). These results mean that Notch activation occurs generally near the DTC/niche nucleus and its proximity determines transcriptional activity. The correlation is even stronger when *sygl-1* mRNA abundance is considered instead of the ATS (Figure 5B), which may be due to the difference in half-lives of *sygl-1* ATS and mRNAs, which is <15 mins or >1 h, respectively (Lee et al. 2016). The correlation between the ATS and the DTC/niche nucleus can be underestimated by a fraction of GSCs that do not exhibit *sygl-1* ATS at the moment of imaging because of the stochastic and sporadic nature of *sygl-1* transcriptional firing (Lee et al. 2016; Lee et al. 2019). In contrast, the retaining mRNAs can be more reflective of transcriptional activation over a longer period, showing the trend more obviously, which provides a useful complementary Notch readout as a metric for Notch activation. Because the DTC/niche nucleus moves away from the distal tip of the gonad during aging, which was also shown in the previous study (Kocsisova et al. 2019), the zone where Notch activation occurs also moves along with the DTC/niche nucleus, causing the age-dependent shift of the Notch-responsive GSC pool (Figure 4).

The DTC/niche nucleus is located at the distal tip of the gonad throughout *C. elegans* development and at YA, when the DTC/niche serves as a migratory cell, promoting the gonad migration and the germline tissue polarization (Byrd et al. 2014; Kimble & White 1981). The shift of the DTC/niche nucleus coincides with the DTC/niche losing its leader function (Byrd et al. 2014; Cecchetelli & Cram 2017), suggesting that the cessation of migratory cues may be involved in the initiation of the DTC/niche aging process.

The strong correlation between Notch activation and the proximity of the DTC/niche nucleus (Figure 5 and Figure S6) indicates that the signaling transduction occurs mostly near the DTC/niche nucleus, implying that most transcripts or proteins of Notch ligands do not travel far but stay close to the nucleus, or that the ligands are more prone to be degraded when they are away from the nucleus. Regardless, the DTC/niche nucleus determines the location of Notch activation in the germline and therefore defines the GSC pool.

### The link between the DTC/niche aging and its structure and morphology

In addition to a shift of its nucleus, the DTC/niche undergoes structural and morphological changes during aging (Figure 6). The DTC processes, including the small intercalating processes (SIPs) and the long extended processes (LEPs), grow longer during aging (Crittenden et al. 2006) (Figure 6B and Figure S7A) when the progenitor zone (PZ) shortens (Figure S1A). One possibility is that Notch ligands, which are transmembrane proteins, can be diluted and more spaced out on the extended processes of the aged gonads, where the total cellular membrane surface increases, weakening overall Notch activation. In addition, the size of the gaps, discontinuous regions in the DTC/niche cap, increases with aging but the number of gaps remains unchanged (Figure 6C-D), meaning that aging itself does not create more gaps but the existing gaps become larger, possibly by the DTC/niche being stretched due to the extending SIPs and LEPs. The larger gaps in the aged DTC/niche hamper the GSC-DTC/niche interactions, causing a decline in Notch activation in the germline. Consistently, there is a decline both in the clustering of *sygl-1* ATS-containing cells to the DTC/niche nucleus (Figure 4D) and the correlation between Notch activation and the proximity of the DTC/niche nucleus (Figure 5) in the older gonads (Figure 5). Notably, the number of the DTC/niche fragments, small fragments detached from the main DTC/niche body, decreases throughout aging (Figure 6E). The small fragments are the result of new branch formation attempts (Wong et al. 2013), thus a reduction of fragments indicates a less active DTC/niche. A decrease in the DTC/niche dynamics also means fewer GSC-DTC/niche interactions, leading to a GSC decline. Therefore, the DTC/niche dynamics can play a role in the aging process, which can be monitored by live imaging with a fluorescence marker for the membrane of the DTC/niche.

## Materials and methods

### Nematode maintenance

All strains were maintained at 20°C as described in (Brenner, 1974). The wild type was N2 Bristol. The transgene was as follows: LGIII: qSi153[Plag-2::myr-GFP; Pttx-3::DsRED] III (Byrd et al. 2014), where the strain was derived). For smFISH and immunofluorescence of the wild type and transgenic stains, animals were grown at 20°C until plates contained adult worms and eggs were present on the agar plate. The animals were synchronized using the hypochlorite treatment or the bleaching technique as described in (Porta-de-la-Riva et al. 2012) to produce synchronized L1 larvae. The L1 larvae continue to grow at 20°C until 24 h post-mid-L4 stage (young adult (YA)). For aged worms (A24-A72), worms were transferred to a new plate every 24 h until 96 h post-mid-L4.

### Strains used in this study

N2: wild type

JK4533: *qSi153[Plag-2::myr-GFP; Pttx-3::DsRED] III*

### Antibodies

Anti-Green Fluorescent Protein Mouse IgG 2A (Fischer Scientific A11120)

IgG (H+L) Highly Cross-Adsorbed Donkey anti-Mouse, Alexa Fluor® Plus 488, Invitrogen™ (Fischer Scientific PIA32766)

### Single Molecule RNA Fluorescent in situ Hybridization and Immunofluorescence (Co smFISH-IF)

Benchtop, gloves, and pipettors were wiped down with RNAseZap and RNase-free filtered tips and tubes were prepared. Synchronized plates were washed with 2 mL of non-RNase free RNase-free 1X Phosphate Buffered Saline solution with 0.1% Tween-20 (PBST) and transferred to a 60mm plastic petri dish cover. Then an additional 2-3 mL of PBST was added to the dish cover. To extrude the gonads, 0.25 M levamisole stock was first added in a 1:1000 dilution to the PBST, attaining a final levamisole concentration of 0.25 mM. Then using a sterilized scalpel, the worms were dissected behind the pharynx or before the rectum to release the gonads. Dissections were completed within 20 minutes to prevent any gonad deformation. Dissected worms were collected in a 1.5 mL Eppendorf tube and spun down for 60 secs at 2000rpm. The supernatant was removed, and the sample was fixed with PBST and 3.7% formaldehyde and incubated on a rotator at room temperature for 30 mins. The sample was then spun down to remove the supernatant into an organic waste bottle. To permeabilize the sample, 1 mL of RNase-free 1X PBS + 0.1% Triton X-100 was added to the sample and incubated on a rotator for 10 mins at room temperature. The sample was spun down at 2000 rpm for 60 secs and the supernatant was removed. The sample was washed twice with 1 mL RNase-free PBST. The sample was spun down at 2000 rpm for 60 secs and was inverted 5-6 times between each wash. The sample was spun down after the second wash and resuspended in 1 mL of 70% of ethanol. Samples were placed at 4°C overnight or up to 1 week.

To prepare the probe for the hybridization step, the dried probe mix (5 nmol) to 40μL of RNase-free TE buffer (10 mM Tris-HCl, 1 mM EDTA, pH 8.0) to create a probe stock of 125 μM. The probe was then diluted to 1:20 (6.25 μM). Fixed samples were spun down and ethanol was removed. Samples were equilibrated in smFISH wash buffer (2X SSC, 10% deionized formamide in nuclease-free water) for 5 mins. While the samples equilibrate in the wash buffer, 48μL of hybridization buffer (HB; Contains 1 g dextran sulfate, 1 mL 20X SSC, 7.3 mL H20 (or up to 10 mL volume), and 1 mL formamide is thawed and added to a new 1.5 mL RNase free tube with 1 μL of each probe dilution (1:20; intron and exon of *sygl-1*; intron probe for *let-858*). The samples were then spun down for 60 secs at 2000 rpm and the supernatants were discarded. The probe-HB mix was then added to the sample and incubated at 37°C for 4-72 h.

After hybridization, 1 mL wash buffer was added, spun down, and removed. To prepare the sample for immunofluorescence, 100 μL of blocking solution (PBST + 0.5% BSA) was added and incubated for 30 mins at room temp. The blocking solution was spun down and removed. The primary antibody was diluted 1:200 in blocking solution and incubated overnight at 4°C. Samples were washed twice with 200 μL of blocking solution. The secondary antibody was diluted 1:1000 in blocking solution and DAPI was added at a concentration of 0.5 μg/mL. The sample was incubated in secondary antibody/DAPI at room temperature for 1-2 h. The sample was washed twice with 200 μL blocking solution, spun down, and removed. The samples were then resuspended in 10 μL of ProLong Gold mounting media and mounted on a glass slide in a dropwise manner. Slides are cured for 24-60 h and sealed.

### Nuclear morphology

DAPI staining was used to assess cell row counts and progression into gametogenesis (mitotic or meiotic cells). Mitotic cells are identified by their round ‘donut-like’ shape due to the lack of DAPI expression in the center from a large nucleolus. Meiotic cells are presented in two manners, crescent-shaped transitionary cells, or as larger rounded cells with visible chromatin condensation in the center (Hillers et al. 2017).

### Distal tip cell nucleus

The DTC/niche nucleus position was identified using DAPI staining. Key morphological differences between the DTC/niche nucleus and the GCSs include a lack of a dark center and an elliptical shape, allowing it to be easily distinguished from the rounded germ cell nuclei (Hillers et al. 2017). For measuring the DTC/niche nuclear shift, the Euclidean distance was measured from the most distal point of the gonad to the center of the DTC/niche nucleus.

### Progenitor zone extent

The progenitor zone extent was assessed by measuring from the most distal end of the gonad to the end of the progenitor zone using a segmented line on ImageJ to consider the natural curve of the gonad. The boundary of the progenitor zone was determined by the presence of more than one cell presenting a crescent-shaped morphology (Crittenden et al. 2006; Byrd et al. 2014).

### DTC processes and cap structure assessment

DTC/niche processes were measured in fixed germlines from the strain JK4533 containing *qSi153(Plag-2::myr-GFP*) stained with anti-GFP. The extent of long external processes (LEPs) was measured from the most distal end of the DTC/niche cap to the most proximal point of a continuous process. The extent of the short intercalating processes (SIPs) was measured from the most distal end of the DTC/niche cap to the most proximal point of the web-like processes. Structural changes to the cap such as gaps were measured using the segmented line function in ImageJ to measure the length of regions of the DTC cap lacking GFP expression as gaps as well as the number of gap occurrences. Detached DTC fragments were identified as GFP blobs > 0.5 μm and are not connected to a process. Due to its amorphous shape, the fragment lengths were measured longwise using a straight-line tool on ImageJ.

### Widefield Microscopy setup with Thunder Processing and image acquisition

Gonads were imaged using a Leica DMi8 (Widefield Microscope) equipped with a Leica HC PL APO 63x/1.40-0.60 NA oil immersion objective, LED8 fluorescence illuminator, and THUNDER Imager with exceptional computational clearing methods to remove excessive background. All gonads were imaged completely (depth >15 μm) with a Z-step size of 0.3 μm using the Leica Application Suite X (LAS X) acquisition software (Leica Microsystems Inc., Buffalo Grove, IL). All imaging was done with LED8 light sources. Channels were sequentially imaged in decreasing wavelengths to avoid bleed-through and prevent any photobleaching from occurring.

The *sygl-1* exon probe (TAMRA) was excited at 555nm (40%), and the signal was acquired at 540-640 nm (gain was set to high well capacity) with an exposure time of 250 ms. The *sygl-1* intron probe (Quasar 670) was excited at 635 nm (40%) and the signal was acquired at 625-775 nm (gain was set to high well capacity) with an exposure time of 250 ms. The *let-858* intron probe (TAMRA) was excited at 555nm (40% illumination), and the signal was acquired at 540-640 nm (gain was set to high well capacity) with an exposure time of 250 ms. GFP was excited at 475 nm (10% illumination), and the signal was acquired at 470-550 nm (gain was set to high well capacity) with an exposure time of 200 ms. DAPI was excited at 390 nm (10% illumination), and the signal was acquired at 400-480 nm (gain high well capacity) with an exposure time of 50ms.

### MATLAB Analysis of Acquired images

All processes were implemented and automated using custom MATLAB codes similar to the source code (developed in our previous work (Lee et al. 2016) with certain modifications to optimize the source code for use with Widefield microscopy images. MATLAB R2021a with the ‘image processing’ toolbox was used for all image processing and analyses. The images acquired in this study were packaged in LIF format and were prepared for MATLAB processing: (1) rotate the image to orient the distal end to the left-hand side (2) cropping to the 54-65 μm mark from the distal end using ImageJ (3) and saving as TIFF format with their metadata (e.g., pixel size, z-step size, and the total number of z slices) intact.

The TIFF files of the images were then read into the MATLAB detection code and the X, Y, and Z coordinates and signal intensity of gonadal boundaries, nuclei, mRNAs, and ATS (nuclear spots) were recorded. Once the channels are separated and recorded, the gonadal boundary is set and determined using an RNA channel. If the image was oriented incorrectly, then the MATLAB code is designed to reorient the distal end to the left for proper further analysis. The MATLAB code reconstructs the gonads in a cylindrical manner with the radius changing throughout the distal-proximal axis. This boundary was then used to eliminate any non-specific fluorescent signals located on the periphery of the gonad. The nuclei were then reconstructed with the DAPI signal in each Z-plane. The MATLAB function ‘imfindcircles’ was used to define each nuclear boundary by drawing concentric nuclear circles. The DAPI signal in each Z-plane was then normalized to the mean background intensity within the same Z-plane, where there were no nuclei present. These concentric circles in different Z-planes were fit together in the process of creating a 3-D spherical nucleus by drawing a best-fit sphere and were scored as a part of the nuclei based on two criteria: (1) concentric circles were detected in ≥4 consecutive Z-planes and (2) the variation in Euclidean distances (X, Y direction) of the centers within the concentric circles was <0.5μm. To estimate the nuclear size and DAPI intensity, the radius of the 3-D spherical nucleus was recorded and the DAPI signal within the 3-D spherical nucleus was summed and recorded, respectively.

For RNA detection, it was divided up into two categories: ATS detection and mRNA detection. For the detection of ATS, a Gaussian local peak detection method (‘FastPeakFind’ function) and a signal-to-local background ratio method were used independently to detect ATS using the channel obtained from the intron-specific probe. The ATS considered in the further analysis were detected in both methods as an ATS. If the ATS was not detected via both methods, the detected ATS was then discarded. Each potential ATS candidate within a Z-plane were detected based on the following criteria for *sygl-1*: (1) the signal-to-mean overall background ratio is greater than 1.0 in the germline; (2) the signal to local background (the area located within 3X the distance from the ATS center) ratio is greater than 1.05; (3) a 1*10^−10^ p-value in t-tests comparing all pixel values in candidate ATS to its local background; and (4) size of the ATS greater than 3×3 pixels. These ATS candidates were then overlaid with the MATLAB-generated nuclei and filtered out for true ATS using the following criteria: 1) RNA spots in the intron-specific smFISH channel were localized to a nucleus; (2) the intron probe nuclear RNA spot is co-localized with its corresponding exon-specific probe nuclear spot; and (3) the signal intensities from both exon and intron probes were at least as bright as a single mRNA spot that is detected.

For the detection of mRNA, the MATLAB code was set up in a similar manner as the ATS detection described above with the exception that it was based on *sygl-1* exon-specific probes. First, the RNA spot localized to the nuclei were detected with exon-specific probes and were removed. mRNA candidates were identified in each Z-plane using the Gaussian local peak detection method described above. The number and intensity of mRNA spots were determined using the size, volume, shape, and fluorescence signal intensity of each mRNA spot. The mean mRNA intensity and the mean mRNA size were then determined for each Z-plane. Next, the 3-D reconstructed gonad was used to remove any spots that were outside of the gonad boundary and were <20% of the mean mRNA intensity and the mean mRNA size. Once the signal intensities for each gonad were determined, these intensities were normalized using the data generated from the exon probes, which are specific to both ATS and mRNAs, across the different germlines and different senescent stages for proper direct comparison. The ATS intensities measured from the exon probes were also used in the direct comparison between ATS and mRNAs intensities and numbers. The mean of cytoplasmic mRNA intensities for each 3-D gonad was set to 1 a.u., and the germ cell boundary was determined using a 3-D Voronoi diagram to estimate the number of mRNAs per germ cell (Ledoux, 2007; Yan et al., 2010). This allows the reconstruction of the germ cell boundary at the midpoint between two neighboring nuclei. The size or radius of each Germ cell was restricted to 3μm from the nucleus center.

After the analysis was completed, MATLAB and GraphPad Prism were used to conduct statistical tests and visualize data where the dataset underwent normality tests (Anderson-Darling Normality test). If the data set met the requirements for parametric statistical analysis, ANOVA and t-tests were performed. If the data set did not satisfy the requirements for parametric analysis, the Kolmogorov-Smirnov (KS) test (a nonparametric version of the t-test) was used to compare data.

## Figure legends

**Figure S1**

(a) The size of the progenitor zone (PZ), the distance between the distal tip to the first cells in meiotic prophase. n=45.

(b) The total number of *sygl-1* ATS in the gonad. n=27, 25, 26, and 24 gonads for YA, A24, A48, and A72, respectively.

(c) The histogram of the summed *sygl-1* ATS intensities. n=621, 460, 508, and 444 gonads for YA, A24, A48, and A72, respectively.

**Figure S2**

(a) Images with smFISH using an intron specific probe set targeting a Notch-independent gene, *let-858* for all ages, from YA to A72. Asterisk: the DTC nucleus. scale bar: 10 μm.

(b-d) The total number of *let-858* ATS (b), the total cell count (c), or the number of let-858 ATS per cell

(d) within 60 μm from the distal end of the gonad. n=10 gonads.

**Figure S3**

(a) The percentage of cells containing at least 1 *sygl-1* ATS as a function of the distance from the distal end. Each bar is also separated by the number of *sygl-1* ATS per cell, from 1 to 4. n=27, 25, 26, and 24 gonads for YA, A24, A48, and A72, respectively.

(b) DAPI staining at the distal gonad in A72. The arrow indicates nuclei at meiotic prophase or at pachytene, morphologically distinct from the miotic nuclei, an indication of premature germ cell differentiation. A single Z plane is shown. Asterisk: the DTC/niche nucleus. Scale bar: 10 μm.

(c) The individual *sygl-1* ATS intensities are plotted against the distance from the distal end in all ages. n=931, 661, 660, and 550 gonads for YA, A24, A48, and A72, respectively.

**Figure S4**

(a) The total number of *sygl-1* mRNAs per cell as a function of distance from the distal end.

(b) The percentage of cells containing *sygl-1* mRNAs above the basal level, which is the mRNA count at 40-60 μm from the distal end, where sygl-1 is not actively transcribed by Notch signaling (see Materials and methods).

(a-b) n=27, 25, 26, and 24 gonads for YA, A24, A48, and A72, respectively.

**Figure S5**

(a) A single Z plane of sygl-1 smFISH images taken with YA or A72 gonads. The arrowhead indicates the location of the DTC/niche nucleus and the enrichment of *sygl-1* transcription, both the nascent RNAs and mRNAs.

(b) The distance between the cells containing *sygl-1* mRNAs above the basal level and the DTC/niche nucleus.

(c) The nearest neighbor distance (NND) of cells with *sygl-1* mRNAs above the basal level.

(d) The size of the GSC pool, estimated by the distance between the distal end and *sygl-1* ATS-containing cell located most proximally.

(b-d) n=27, 25, 26, and 24 gonads for YA, A24, A48, and A72, respectively.

(e) The size of the *sygl-1* mRNA-rich area. n=45 for all ages.

**Figure S6**

(a-b) The summed *sygl-1* ATS intensities in each cell (a) or the number of *sygl-1* mRNAs per cell (b) from one gonad are plotted against the distance between the corresponding cell and the DTC/niche nucleus. ‘r’ indicates the Pearson’s correlation coefficient. The blue dashed line indicates the line fitting model to calculate Pearson’s r value.

**Figure S7**

(a) The length of the long external process (LEP) is measured in all ages. n=30, 30, 30, and 29 gonads for YA, A24, A48, and A72, respectively.

(b) A single Z plane of the myr-GFP image shown in Figure 6A for each age. Arrow: the DTC gap. Asterisk: the DTC/niche nucleus.

(c) The length of the detached DTC fragments. n=10 for all ages.

