## Supplementary figures and images for "Age-dependent structural and morphological changes of the stem cell niche disrupt spatiotemporal regulation of stem cells and drive tissue disintegration"

### Supplementary figures

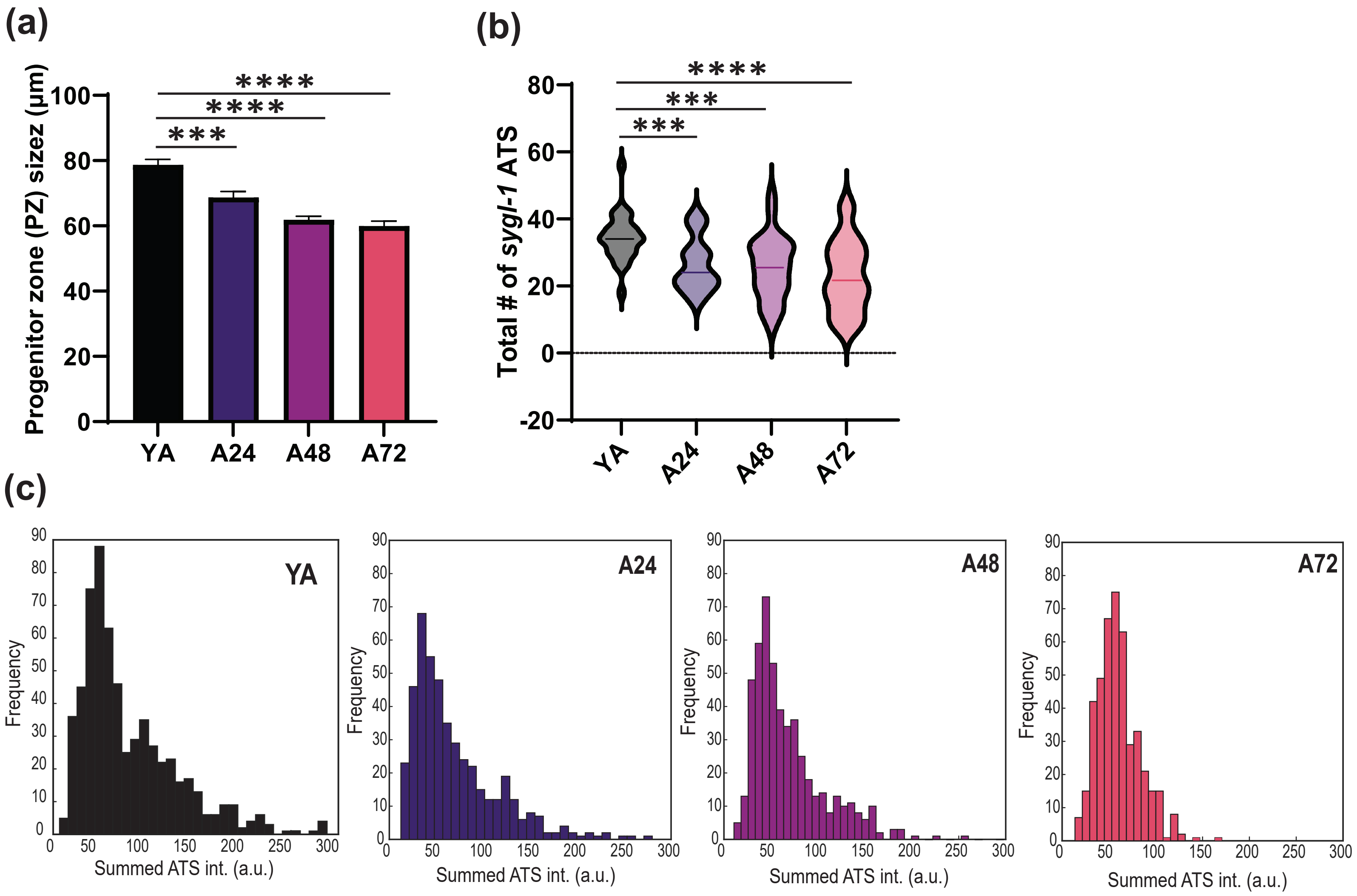

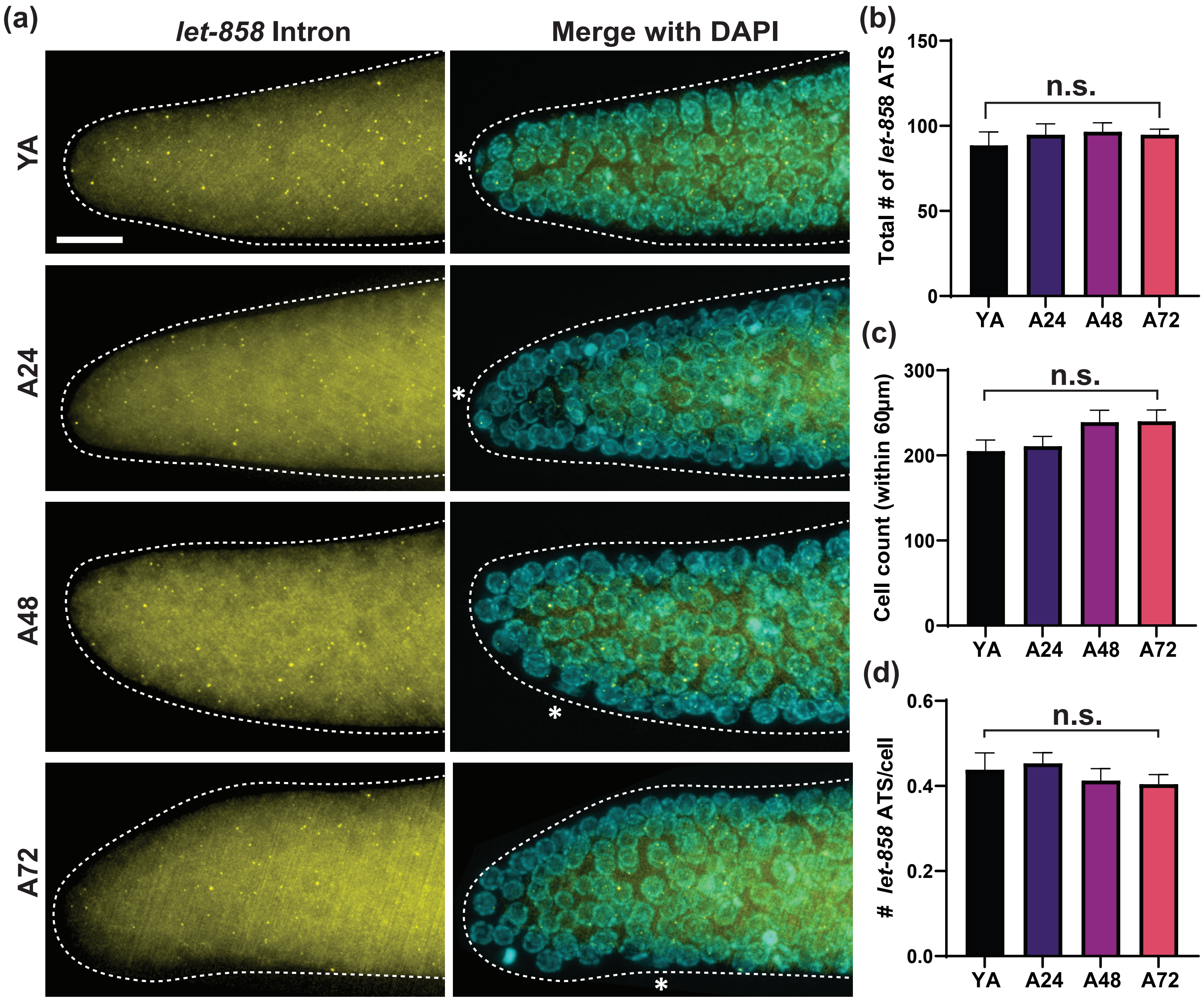

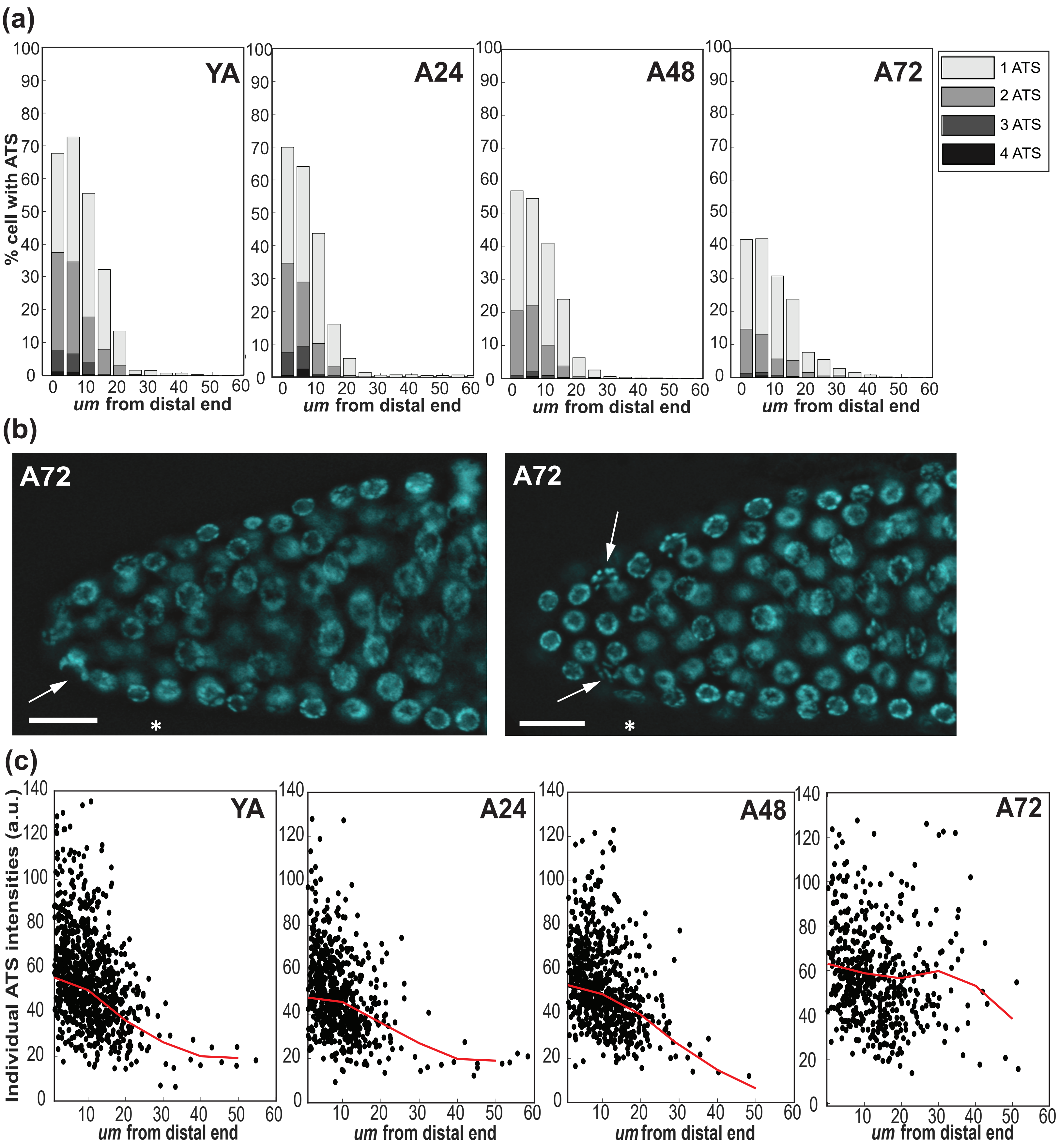

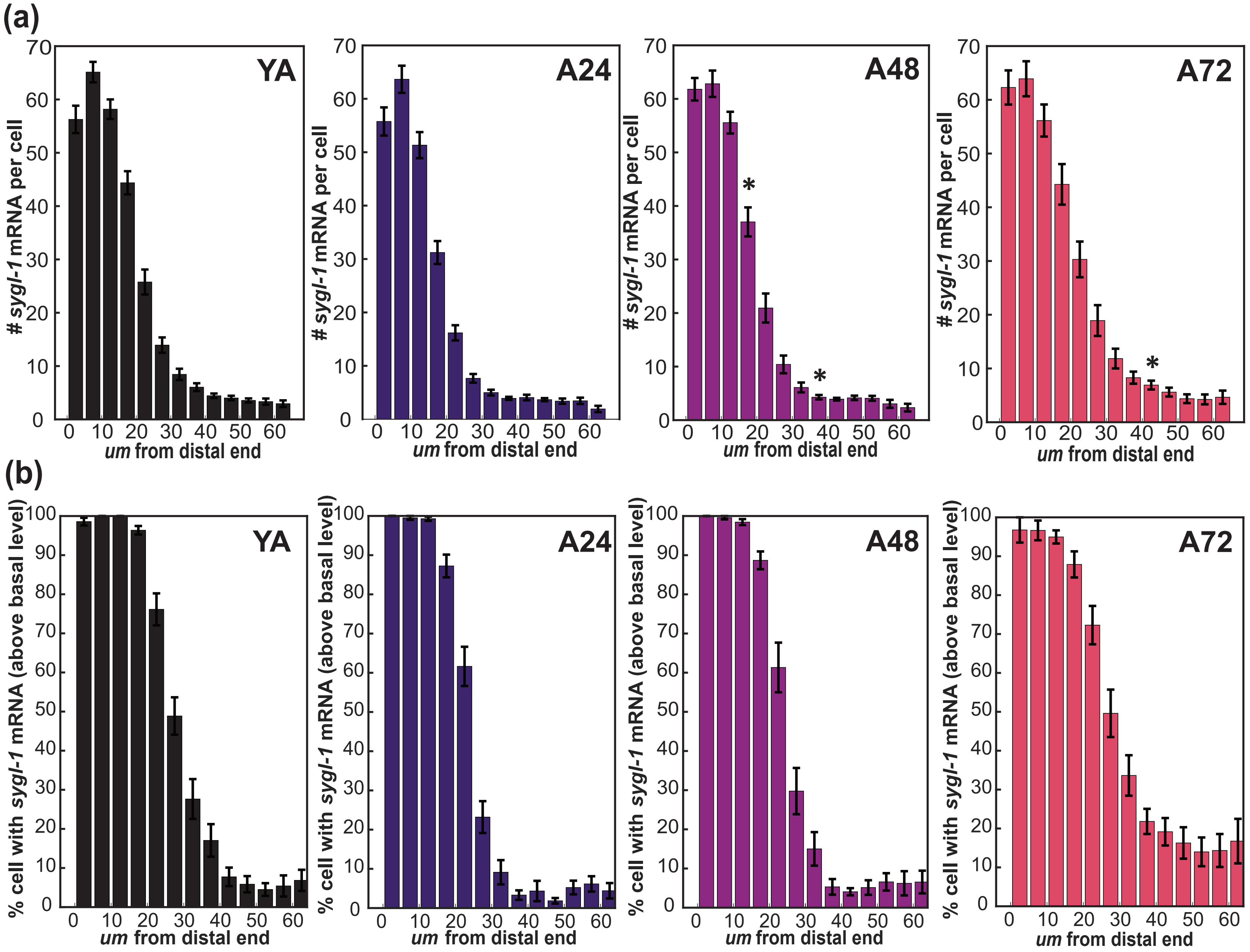

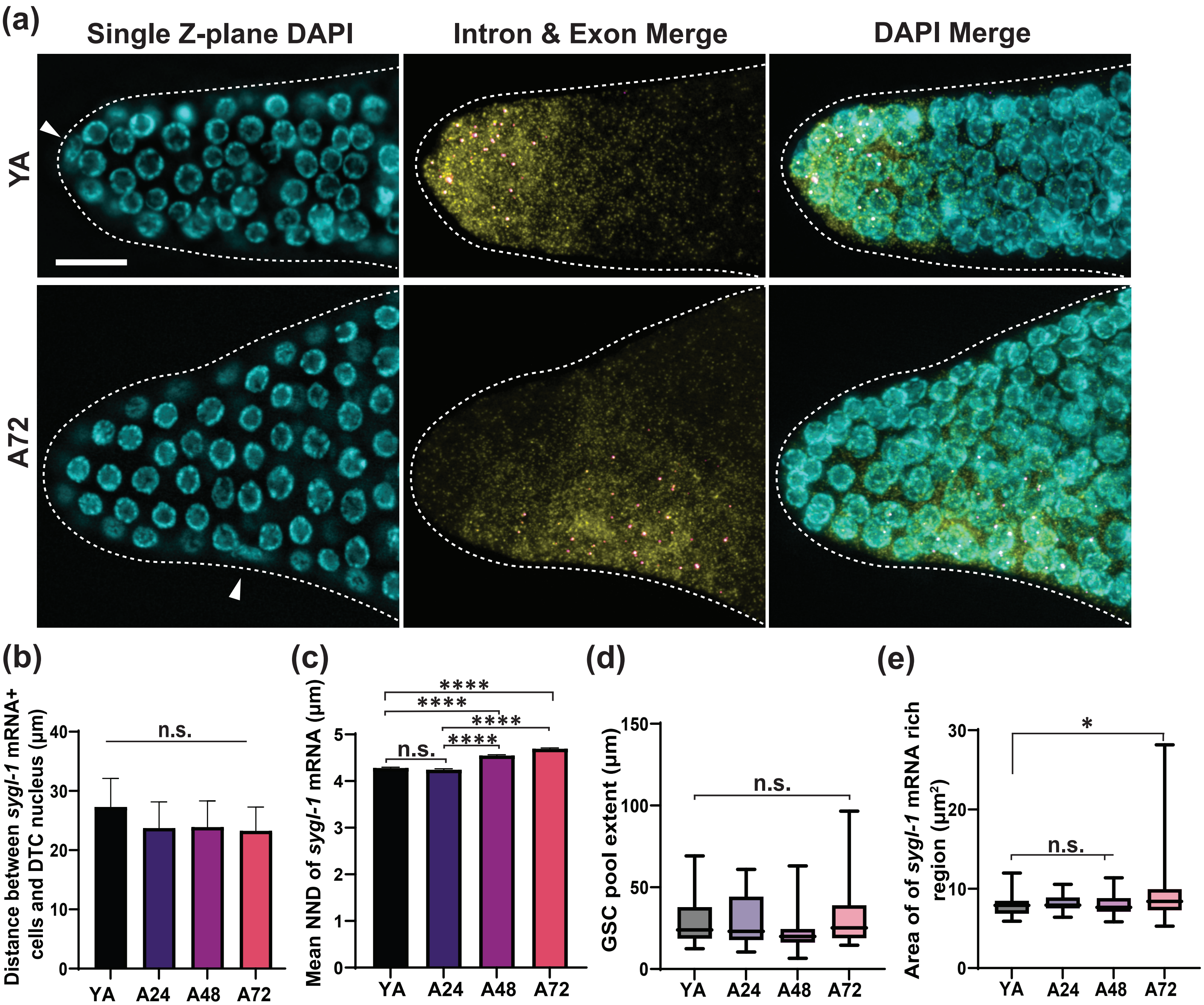

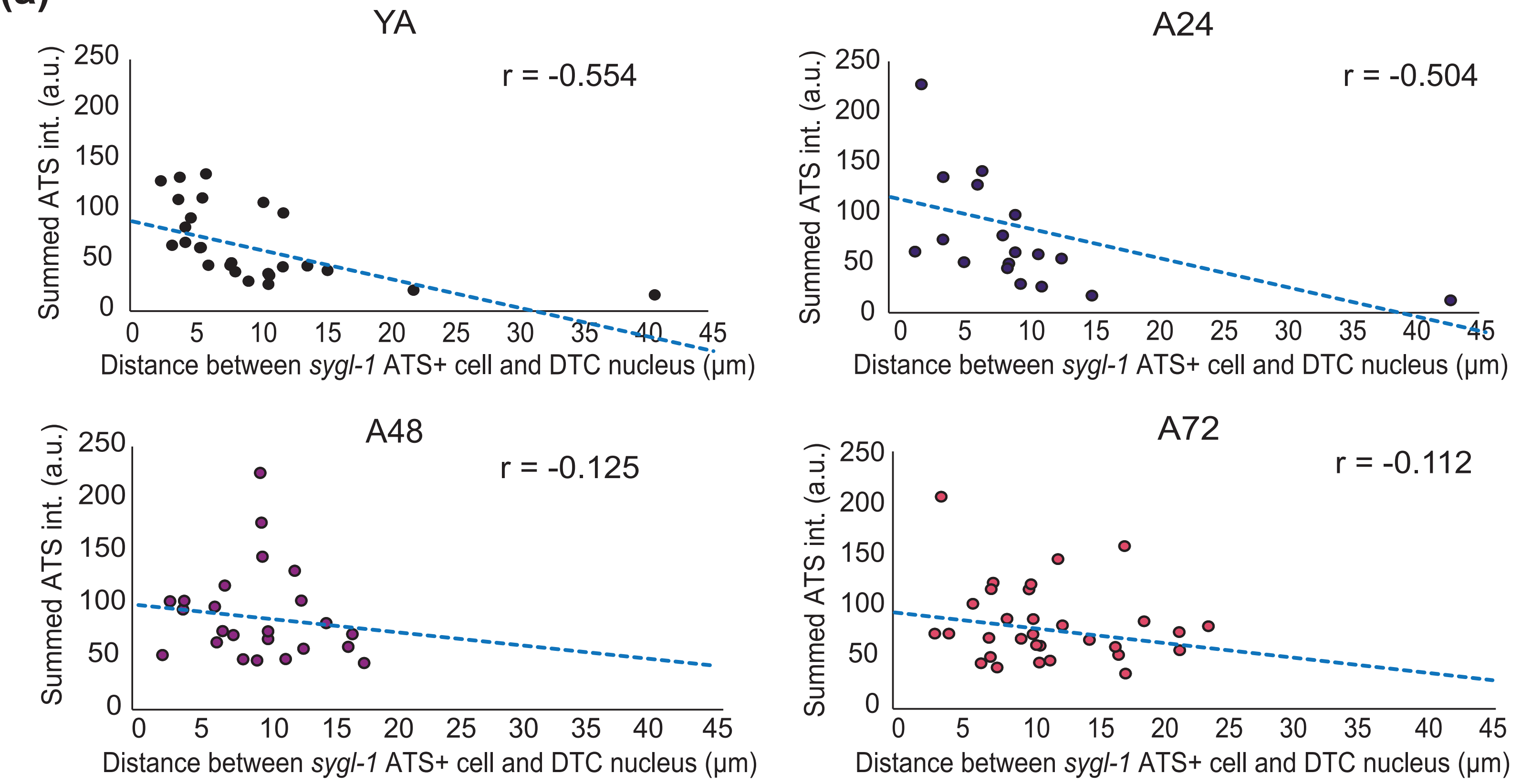

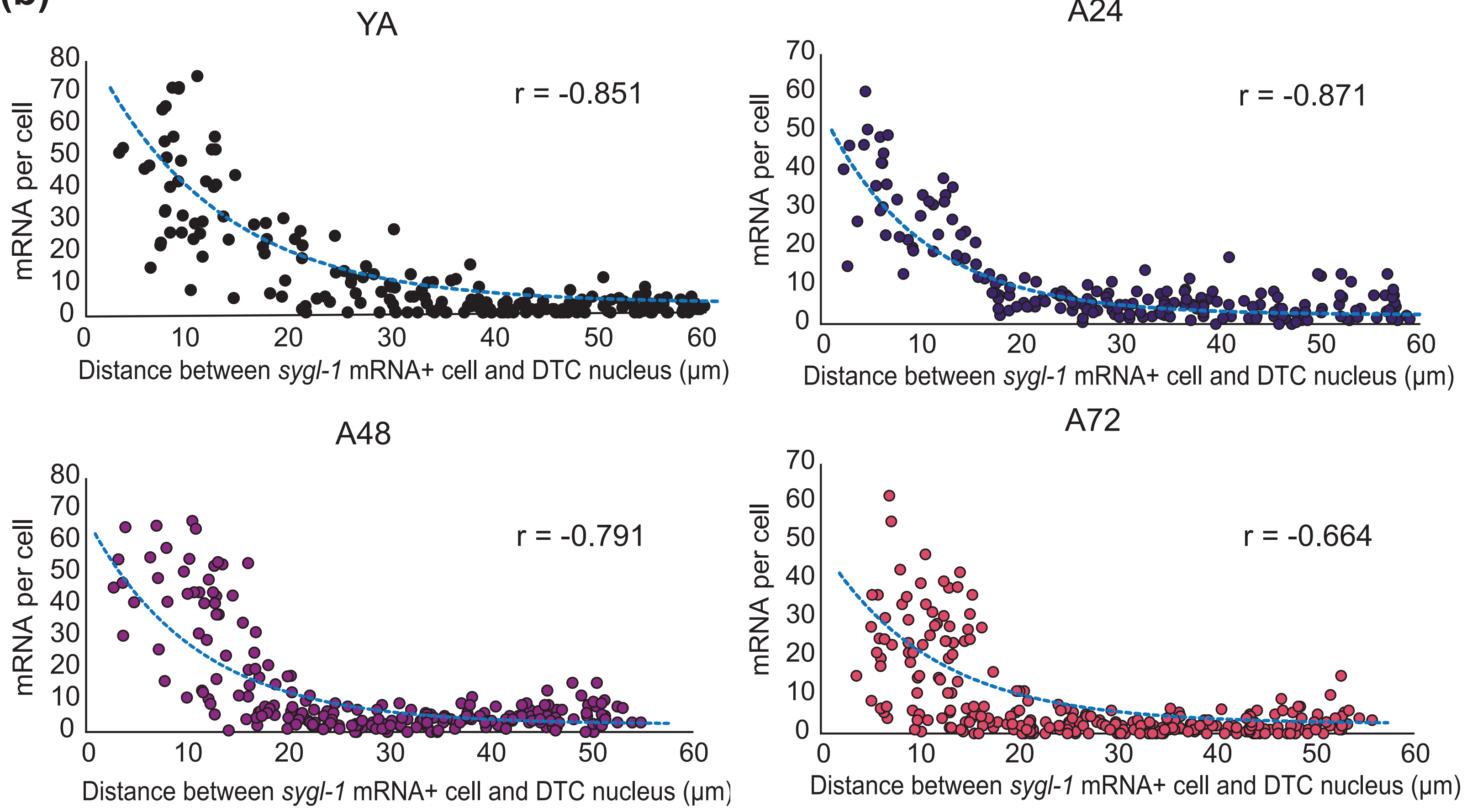

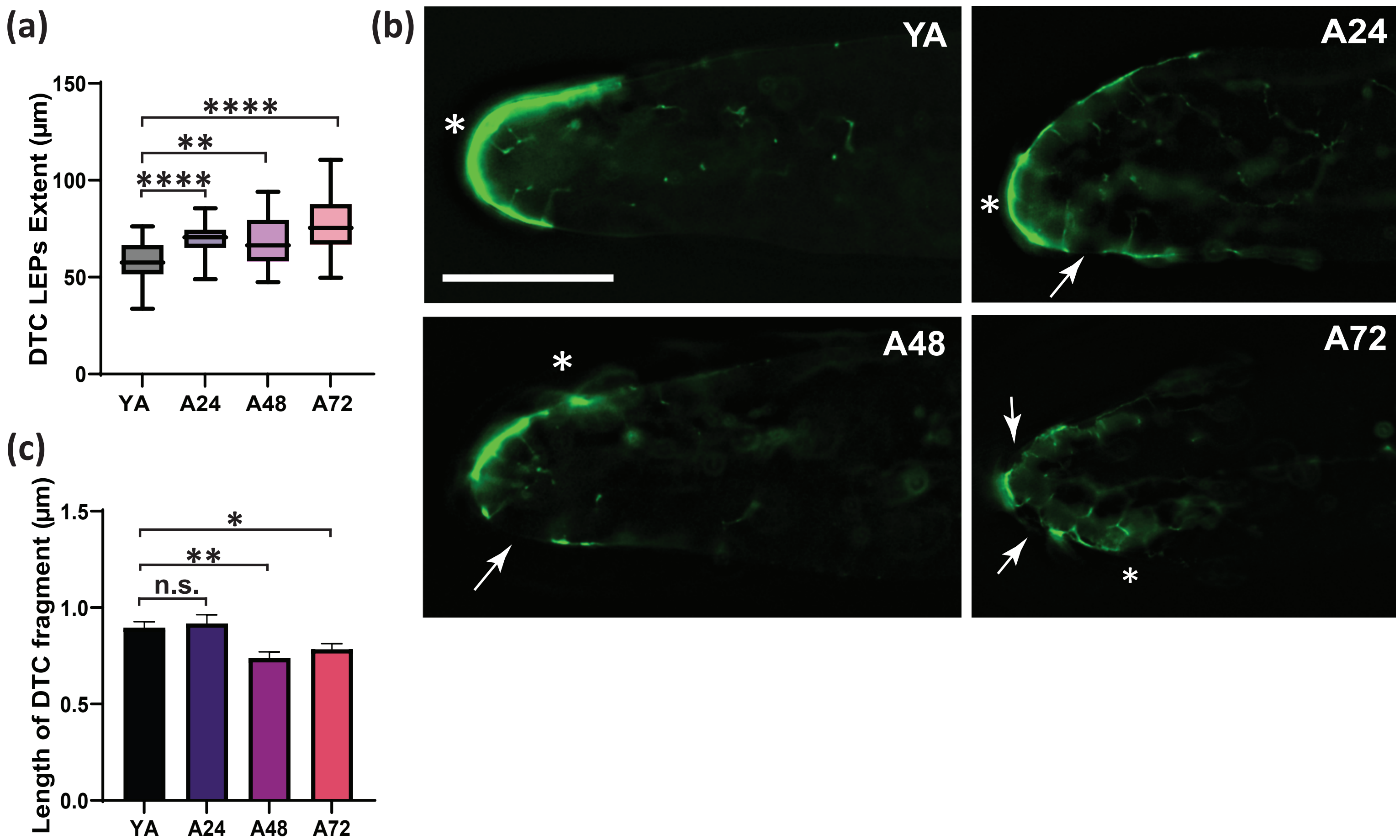
